# Spiking neural network models of sound localisation via a massively collaborative process

**DOI:** 10.1101/2024.07.19.604252

**Authors:** Marcus Ghosh, Karim G. Habashy, Francesco De Santis, Tomas Fiers, Dilay Fidan Erçelik, Balázs Mészáros, Zachary Friedenberger, Gabriel Béna, Mingxuan Hong, Umar Abubacar, Rory T. Byrne, Juan Luis Riquelme, Yuhan Helena Liu, Ido Aizenbud, Brendan A. Bicknell, Volker Bormuth, Alberto Antonietti, Dan F. M. Goodman

## Abstract

Neuroscientists are increasingly initiating large-scale collaborations which bring together tens to hundreds of researchers. However, while these projects represent a step-change in scale, they retain a traditional structure with centralised funding, participating laboratories and data sharing on publication. Inspired by an open-source project in pure mathematics, we set out to test the feasibility of an alternative structure by running a grassroots, massively collaborative project in computational neuroscience. To do so, we launched a public Git repository, with code for training spiking neural networks to solve a sound localisation task via surrogate gradient descent. We then invited anyone, anywhere to use this code as a springboard for exploring questions of interest to them, and encouraged participants to share their work both asynchronously through Git and synchronously at monthly online workshops. At a scientific level, our work investigated how a range of biologically-relevant parameters, from time delays to membrane time constants and levels of inhibition, could impact sound localisation in networks of spiking units. At a more macro-level, our project brought together 31 researchers from multiple countries, provided hands-on research experience to early career participants, and opportunities for supervision and teaching to later career participants. Looking ahead, our project provides a glimpse of what open, collaborative science could look like and provides a necessary, tentative step towards it.

## 1. Introduction

Inspired by the success of endeavours like the Human Genome Project and CERN, neuroscientists are increasingly initiating large-scale collaborations. Though, how to best structure these projects remains an open-question (Mainen et al., 2016). The largest efforts, e.g. the International Brain Laboratory (Abbott et al., 2017; Wool, 2020), The Blue Brain Project and Human Brain Project bring together tens to hundreds of researchers across multiple laboratories. However, while these projects represent a step-change in scale, they retain a legacy structure which resembles a consortia grant. I.e. there are participating laboratories who collaborate together and then make their data, methods and results available upon publication. As such, interested participants face a high barrier to entry: joining a participating laboratory, initiating a collaboration with the project, or awaiting publications. So how could these projects be structured differently?

One alternative is a bench marking contest, in which participants compete to obtain the best score on a specific task. Such contests have driven progress in fields from machine learning (Deng et al., 2009) to protein folding, and have begun to enter neuroscience. For example, in Brain-Score (Schrimpf et al., 2018; 2020) participants submit models, capable of completing a visual processing task, which are then ranked according to a quantitative metric. As participants can compete both remotely and independently, these contests offer a significantly lower barrier to entry. Though, they emphasise competition over collaboration, and critically they require a well defined, quantifiable endpoint. In Brain-Score, this endpoint is a composite metric which describes the model’s similarity to experimental data in terms of both behaviour and unit activity. However, this metric’s relevance is debatable (Bowers et al., 2022) and more broadly, defining clear endpoints for neuroscientific questions remains challenging.

Another alternative is massively collaborative projects in which participants work together to solve a common goal. For example, in the Polymath Project unsolved mathematical problems are posed, and then participants share comments, ideas and equations online as they collectively work towards solutions. Similarly, the Busy Beaver Challenge recently announced a formal proof of a conjecture that was open for decades, based mainly on contributions from amateur mathematicians, organised purely online. Inspired by this approach, we founded COMOB (Collaborative Modelling of the Brain) - an open-source movement, which aims to tackle neuroscientific questions. Here, we share our experiences and results from our first project, in which we explored spiking neural network models of sound localization.

We start by detailing how we ran the project in terms of organisation and infrastructure in Section 2. We then briefly summarise our scientific results in Section 3. We conclude the main text with a Section 4 of what went well, what went wrong, and how we think future projects of this sort could learn from our experiences. Finally, in the Section 7 we provide longer, detailed write-ups of some of our scientific results.

## 2. Towards open collaborative science

### 2.a. Workflow

Our project grew out of a tutorial at the Cosyne conference (2022) for which we provided video lectures and code online (Goodman et al., 2022). Participants joining the project were encouraged to review this material, and then to work through an introductory Jupyter Notebook (Kluyver et al., 2016) containing Python code, figures and markdown text, which taught them how to train a spiking neural network to perform a sound localisation task.

Participants were then directed to our website where we maintained a list of open scientific and technical questions for inspiration. For example, how does the use of different neuron models impact network performance and can we learn input delays with gradient descent? Then, with a proposed or novel question in hand, participants were free to approach their work as they wished. In practice, much like a “typical” research project, most work was conducted individually, shared at monthly online meetings and then iteratively improved upon. For example, several early career researchers tackled questions full-time as their dissertation or thesis work and benefited from external input at monthly workshops.

**Figure 1:**
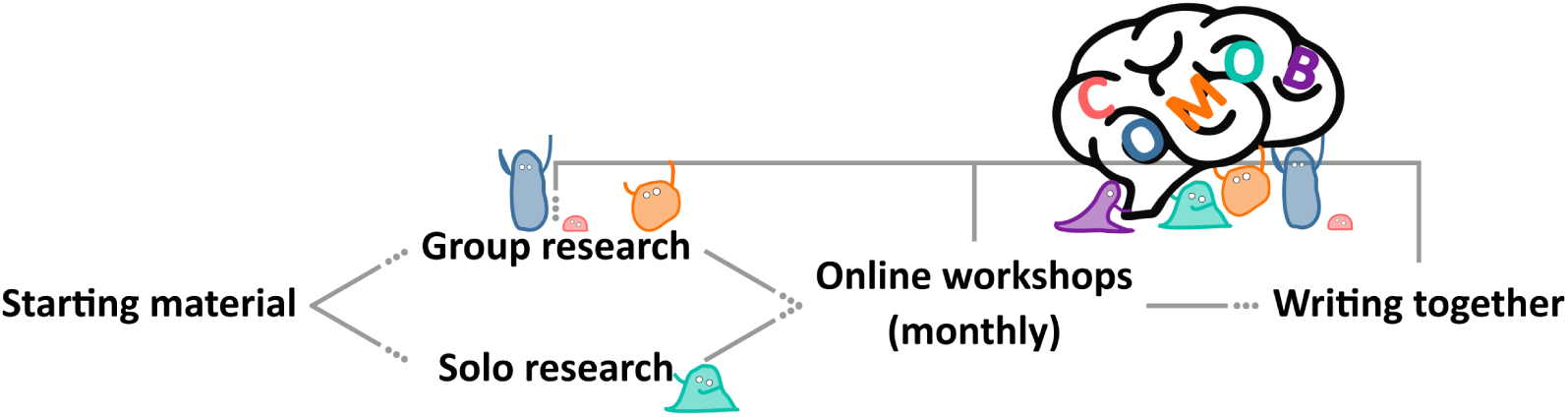
Our workflow. To on-board new participants we provided text, videos and code outlining our scientific starting point. This material formed a springboard for participants to pursue research either individually or in small groups. We then iteratively improved on this work through monthly online workshops and by writing this paper together through an open, collaborative process.

We consciously decided on this free-form structure to experiment with the feasibility of a more bottom-up approach to doing team science, with minimal top-down supervision. We discuss the advantages and disadvantages of this approach in the Section 4.

### 2.b. Infrastructure

#### 2.b.i. Code

We provided a Starting Notebook to give participants an easy way to get started. This is a Python-based Jupyter Notebook (Kluyver et al., 2016), an interactive cell-based environment which allows a mixture of text, image and code cells to be weaved together, combined with an easy user interface. Participants could either work locally using their own Python distribution, or using a cloud compute service such as Google Colab. We choose Google Colab to minimise the entry barrier for participation, as it is a free service (although blocked in certain countries unfortunately) where all the software packages needed are pre-installed, meaning it takes users only a few seconds to go from reading about the project to running the code.

Our starting notebook used a combination of NumPy (Harris et al., 2020), Matplotlib (Hunter, 2007), and PyTorch (Paszke et al., 2019). The code for surrogate gradient descent was based on Friedemann Zenke’s SPyTorch tutorial (Zenke, 2019; Zenke & Ganguli, 2018).

Note that we didn’t require participants to use our starting notebook, and indeed in Section 7.d, De Santis and Antonietti implemented a very different sound localization model from scratch.

#### 2.b.ii. GitHub

Like many open-source efforts, our public GitHub repository was the heart of our project. This provided us with three main benefits. First, it made joining the project as simple as cloning and committing to the repository. Second, it allowed us to collaborate asynchronously. That is, we could easily complete work in our own time, and then share it with the group later. Third, it allowed us to track contributions to the project. Measured in this way, 28 individuals contributed to the project. However, interpreting this number is challenging, as these contributions vary significantly in size, and participants who worked in pairs or small groups, often contributed under a single username. We return to this point in the Section 4.

#### 2.b.iii. Website via MyST Markdown

For those interested in pursuing a similar project our repository can easily be used as a template. It consists of a collection of documents written in Markdown and executable Jupyter Notebooks (Kluyver et al., 2016) containing all the code for the project. Each time the repository is updated, GitHub automatically builds these documents and notebooks into a website so that the current state of the project can be seen by simply navigating to the project website. We used MyST Markdown to automate this process with minimal effort. This paper itself was written using these tools and was publicly visible throughout the project write-up.

### 2.c. Teaching with this framework

This project emerged from a tutorial, and the code remains well suited for teaching several concepts from across neuroscience. As such, we integrated our project into a Physics MSc course on Biophysics and Neural Circuits. Working individually or in pairs, students actively engaged by adjusting network parameters and modifying the provided code to test their own hypotheses. Later, brief progress report presentations stimulated dynamic discussions in class, as all students, while working on the same project and code, pursued different hypotheses. We found that this setup naturally piqued interest in their peers’ presentations, enhanced their understanding of various project applications, and facilitated collaborative learning. Moreover, it allowed for engagement from students at a range of skill levels, and helped bridge the gap between teaching and research. For those interested in teaching with this framework, introductory slides and a dedicated Python notebook are available on our GitHub repository.

## 3. Training SNNs for sound localisation

### 3.a. Introduction

In the Cosyne tutorial (Goodman et al., 2022) on spiking neural networks (SNNs) that launched this project, we used a sound localisation task. Reasoning that sound localisation requires the precise temporal processing of spikes at which these networks would excel.

Animals localise sounds by detecting location- or direction-specific cues in the signals that arrive at their ears. Some of the most important sources of cues (although not the only ones) come from differences in the signals between two ears, including both level and timing differences. Respectively, termed interaural level difference (ILD) and interaural timing difference (ITD). In some cases humans are able to detect arrival time differences as small as 20 *µ*s.

The classic model of ITD sensitivity is the delay line model of Jeffress (1948) in which an array of binaural coincidence detector neurons receive inputs from the two ears with different delays. When a neurons’ delays exactly match the acoustic delays induced by the sound location, it will be maximally active. Therefore, the identity of the most active neuron indicates the direction of the sound. This model is widely accepted, though was shown to be inefficient with respect to neural noise by McAlpine & Grothe (2003), who proposed an alternative model based on the two binaural hemispheres average firing rates - which is optimally robust to neural noise. However, Goodman et al. (2013) showed that these models perform too poorly to account for behavioural data, especially in situations where sounds had complex and unknown spectral properties, or in the presence of background noise, and proposed an alternative based on a perceptron-like neural network - which is both robust to neural noise and performed well across a range of conditions.

Building on this literature, and our Cosyne tutorial, the starting point of this project was to ask: what solutions would you find if you directly optimised a spiking neural network to localise sounds? How would those solutions depend on the available neural mechanisms and statistics of the sound? Could we understand the solutions found in a simple way? What properties would the solution have in terms of robustness to noise, generalisation, and so forth? And could the solutions found by optimisation throw light on features found in the auditory systems of different animals?

### 3.b. A simple spiking neural network model

We started with a simple, spiking neural network model (described in more detail in Section 7.a.i) trained to solve a highly abstracted sound localisation task.

**Figure 2:**
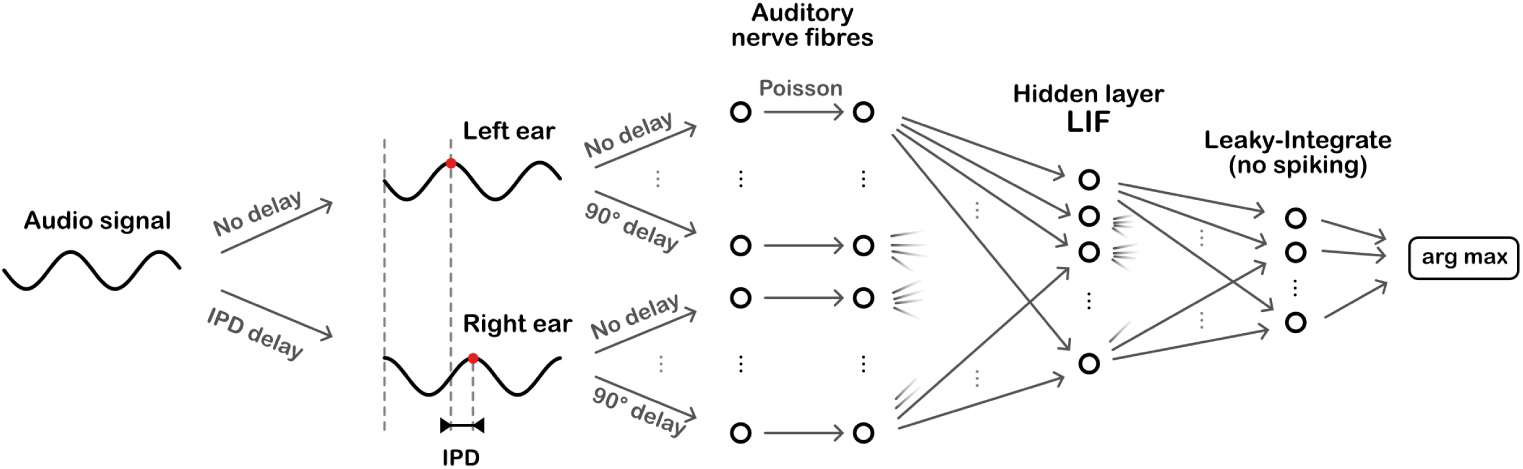
Overall model architecture. Sinusoidal audio signals are passed through two populations of units, with a range of preassigned phase delays, representing the left and right ears. These units generate Poisson spike trains which pass forward to a layer of leaky-integrate and fire (LIF) units, then a layer of leaky-integrator output units from which we readout the networks estimate of the interaural phase difference (IPD).

The task is to estimate the ITD of a pure tone (sine wave) at a fixed frequency. This is equivalent to estimating the interaural phase difference (IPD) since the ITD is ambiguous for a sine wave. The model consists of three layers. First, a layer of spiking input neurons which fire spikes according to a Poisson process with a time-varying rate determined by the input stimulus. This layer is divided into two subpopulations corresponding to the two ears, with signals to one ear delayed with respect to the other. Each neuron within a subpopulation has a different phase delay. Next, comes a single hidden layer of leaky integrate-and-fire (LIF) neurons, and an output layer of leaky, non-spiking neurons. Each output neuron is associated to a particular IPD, and the estimated IPD of the model is the identity of the most active output neuron.

The input neurons are all-to-all connected to the layer of hidden neurons via a trainable weight matrix. In this way, during training the model is free to *select* the neurons with the appropriate phase delays to generate the desired properties for the hidden layer neurons. This lets the model learn to make use of delays without having to directly implement trainable delays, as this is a challenging problem (which we tackled later in Section 3.d).

Using this setup, we successfully trained SNNs on this task, and found that accuracy increased as we reduced the membrane time constant of the units in the hidden layer (Improving Performance: Optimizing the membrane time constant). This initially suggested that coincidence detection played an important role. However, further analysis in../../research/ time-constant-solutions.ipynb (described in more detail in Section 7.a) showed that in fact, the network was not using a coincidence detection strategy, or indeed a spike timing strategy.

Rather, it appears to be using an approach similar to the equalisation-cancellation theory (Culling, 2020; Durlach, 1963) by subtracting various pairs of signals to find the point where they approximately cancel. Careful analysis of the trained model showed that it could be extremely well approximated by a 6-parameter model that is quite easy to describe, but does not obviously correspond to any known features of the auditory system.

As an alternative approach, we also used Tensor Component Analysis (TCA) (Williams et al., 2018) to explore the spiking activity of this model, and to compare it across multiple trained networks (Section 7.b).

Building on this base model, we explored two main questions: how changing the neuron model alters the network’s behaviour and how the phase delays (within each ear) can be learned.

### 3.c. Alternative neuron models

#### 3.c.i. Dale’s principle

In biological networks most neurons release the same set of transmitters from all of their synapses, and so can be broadly be considered to be excitatory or inhibitory to their post-synaptic partners; a phenomenon known as Dale’s principle (Dale, 1935; Strata & Harvey, 1999). In contrast, most neural network models, including our base model, allow single units to have both positive and negative output weights.

To test the impact of restricting units to being either excitatory or inhibitory, we trained our base model across a range of inhibitory:excitatory unit ratios, and tested it’s performance on unseen, test data (Sound localisation following Dale’ law). We found that networks which balanced excitation and inhibition performed significantly better than both inhibition-only networks - which perform at chance level as no spikes propagate forward, and excitation-only networks - which were roughly 30% less accurate than balanced networks.

To understand where in the network inhibition is required, we then trained a second set of networks in which we forced either the input or hidden units to be all excitatory, and set the remaining units to be half inhibitory and half excitatory. Networks with all excitatory hidden units performed as well as networks with balanced units, while networks with purely excitatory inputs performed significantly worse, demonstrating a role for inhibition in the input-hidden connections / delay lines.

Inhibition therefore plays an important role in this model, in line with experimental data that shows that blocking inhibition eliminates ITD-sensitivity in the medial superior olive (Brand et al., 2002; Pecka et al., 2008).

#### 3.c.ii. Filter-and-fire

Unlike most point neuron models, in which pairs are connected by a single weight, many biological neurons make multiple contacts with their post-synaptic partners at different points along their dendritic tree. These contacts evoke post-synaptic potentials (PSPs) with distinct temporal dynamics, depending on their distance from the soma, with distal/proximal contacts inducing prolonged/brief PSPs. These features are captured by the filter-and-fire neuron model (F&F) (Beniaguev et al., 2024), in which units make multiple contacts with their partners and each input is convolved with a distance-from-soma dependent synaptic filter. While networks of F&F units outperform networks of LIF units on a temporal version of MNIST, we hypothesised that this difference would be magnified in our sound localisation task, given it’s natural temporal structure. We found that while training performance was increased using the F&F model, test performance was much worse, suggesting overfitting.

### 3.d. Learning delays

As in our base model, many studies incorporate axonal and/or dendritic delays as non-learnable parameters. Here, we explore how these phase delays, as well as synaptic delays, can be learned through two approaches.

The first method was to develop a differentiable delay layer (DDL). This method uses a combination of translation and interpolation, where the interpolation allows the delays to be differentiable even though time steps are discrete. This can be placed between any two layers in a spiking neural network, and is capable of solving the task without weight training. This work is described in more detail in Section 7.c.

While we were developing our DDL-based method, a paper introducing synaptic delays using dilated convolutions with learnable spacings (DCLS) was published (Hammouamri et al., 2024; Hassani et al., 2023), prompting us to explore this approach as well. This method also relies on interpolation and is very similar to the DDL method, serving as a generalization for synaptic delays. It uses a 1D convolution through time to simulate delays between consecutive layers. The kernels include a single non-zero weight per synapse, which corresponds to the desired delay. This method co-trains weights and delays.

We found that both methods performed well and eliminated the artificial phase delays introduced in the basic model.

### 3.e. Detailed inhibition-based model

Finally, we developed a more detailed model in which we used over 170,000 units, with conductance-based synapses, to approximate the structure of the mammalian brainstem circuit (see more details in Section 7.d).

In short, input spectrograms representing sounds at azimuth angles from −90° to +90° were converted into spikes, then passed forward to populations representing the globular and spherical bushy cells, and subsequently the lateral and medial superior olivary nuclei, from which we readout sound source angle predictions. Note that, unlike the work with our base model, we used no learnable parameters in this model, and instead based parameters on neurophysiological data. For example, the MSO units had excitatory inputs from both the ipsi and contralateral SBCs and dominant inhibition from contralateral GBCs.

This model generated realistic tuning curves for lateral and medial superior olive (LSO and MSO) neurons. Moreover, removing inhibition shifted ITD sensitivity to the midline, as in (Brand et al., 2002; Pecka et al., 2008).

## 4. Discussion

### 4.a. What went well

The decision to start from the code base of the Cosyne tutorial (Goodman et al., 2022) was very helpful. It meant that users had a clear entry path for the project without needing prior expertise, and a code base that was designed to be easy to understand. In addition, the popularity of the tutorial (over 30k views on YouTube at the time of writing) meant that many people heard about this project and were interested in participating. In addition, the GitHub-based infrastructure allowed for asynchronous work and a website that was automatically updated each time anyone made a change to their code or to the text of the paper.

By providing models which used spiking neurons to transform sensory inputs into behavioural outputs, participants were free to explore in virtually any direction they wished, much like an open-world or sandbox video game. Indeed over the course of the project we explored the full sensory-motor transformation from manipulating the nature of the input signals to perturbing unit activity and assessing network behaviour. Consequently, our code forms an excellent basis for teaching, as concepts from across neuroscience can be introduced and then implemented in class. In this direction, we integrated our project into two university courses and provide slides and a highly annotated python notebook, for those interested in teaching with these models.

Beyond providing teaching and hands-on research experience, the project also offered many opportunities for participants to improve their “soft” scientific skills. For early career researchers (undergraduate and master’s students) these included learning how to work with Git, collaborate with researchers from diverse countries and career stages, and contribute to a scientific publication. For later career researchers (PhD, Postdoc) the project provided many supervision and leadership opportunities. For example, during online workshops, later career participants were encouraged to lead small groups focussed on tackling specific questions.

### 4.b. What went wrong

While our sandbox design offered several advantages (discussed above), the open nature of the process did present three challenges. Our first challenge was standardising work across participants; for example, ensuring that everyone used the same code and hyperparameters. Along these lines, future projects would benefit from having participants dedicated to maintaining the code base and standardising participants work.

Our second challenge, was the project’s exploratory nature. While this appealed to many participants, the lack of a clear goal or end-point may have been off-putting to others. For future efforts, one alternative would be to define clear goals *a priori*, however if handled carelessly, this runs the risk of reducing to a to-do list passed from more senior to more junior researchers. A more appealing alternative could be to structure the project in clearly defined phases. For example, early months reviewing literature could be followed by a period of proposals and question refinement, before a final stretch of research. Another alternative, would be to begin by collecting project proposals and allowing participants to vote on a project to pursue. This could be done by having each participant check a box for each project they would choose to work on if it were selected, and then selecting the project with the most checks (thereby maximising the expected number of participants). Multiple projects could even be launched simultaneously if this approach proved popular.

A third challenge, which arose towards the end of the project, was how to fairly assign credit. We had initially - and perhaps somewhat idealistically - stated that anyone who contributed to the project, either by writing code or participating in one of the workshops, would be included on the author list. To the extent that it was possible, we have followed through with this, though we were simply unable to contact several of the participants and so could not include them as authors. Another issue with this system is that participants with unequal contributions, e.g. attending a workshop vs contributing an entire section of the paper, would be assigned similar credit, i.e. authorship. To resolve this, we experimented with using the number or size of GitHub commits to order authors, however we found that these metrics did not accurately reflect contributions. For example, it may be quicker to commit a large-amount of low quality text than a concise well written section, and similarly there is no good reason to distinguish between two authors who submit the same amount of work through a different number of commits. We attempted to address this challenge by providing a contributions table (Section 5) and agreeing an author order. This order was agreed on unanimously, though could easily cause issues in other projects. Consequently, we recommend that a strategy for credit assignment be determined collaboratively at the start of the project, and made explicit so that participants can clearly understand how their contribution will translate to credit. Alternatively, such projects could publish under a pseudonym, e.g. COMOB.

Ultimately, while the project explored many interesting directions, which will form the basis for future work, we did not reach a point where we could draw strong scientific conclusions about sound localization. From group discussions we concluded that this is likely due to the free-form nature of our project, which would have benefited from a more coordinated approach. The question is, how to do this without compromising the ideals of a grass-roots project? Extending the voting idea above, one approach would be to make the proposer of the, democratically selected, project responsible for making sure that results are comparable and generally keeping the project on the right track. A role similar to a traditional supervisor, but with the critical difference that they are elected by their peers and only on a project by project basis.

4.c. *Conclusions*

This paper does not present a scientific breakthrough. However, it does demonstrate the feasibility of open research projects which bring together large number of participants across countries and career stages to work together collaboratively on scientific projects. Moreover, several follow-up research projects are planned based on pilot data from our work and, building on our experience, we plan to launch a second COMOB project soon.

## 5. Contributors

**Table 1:**
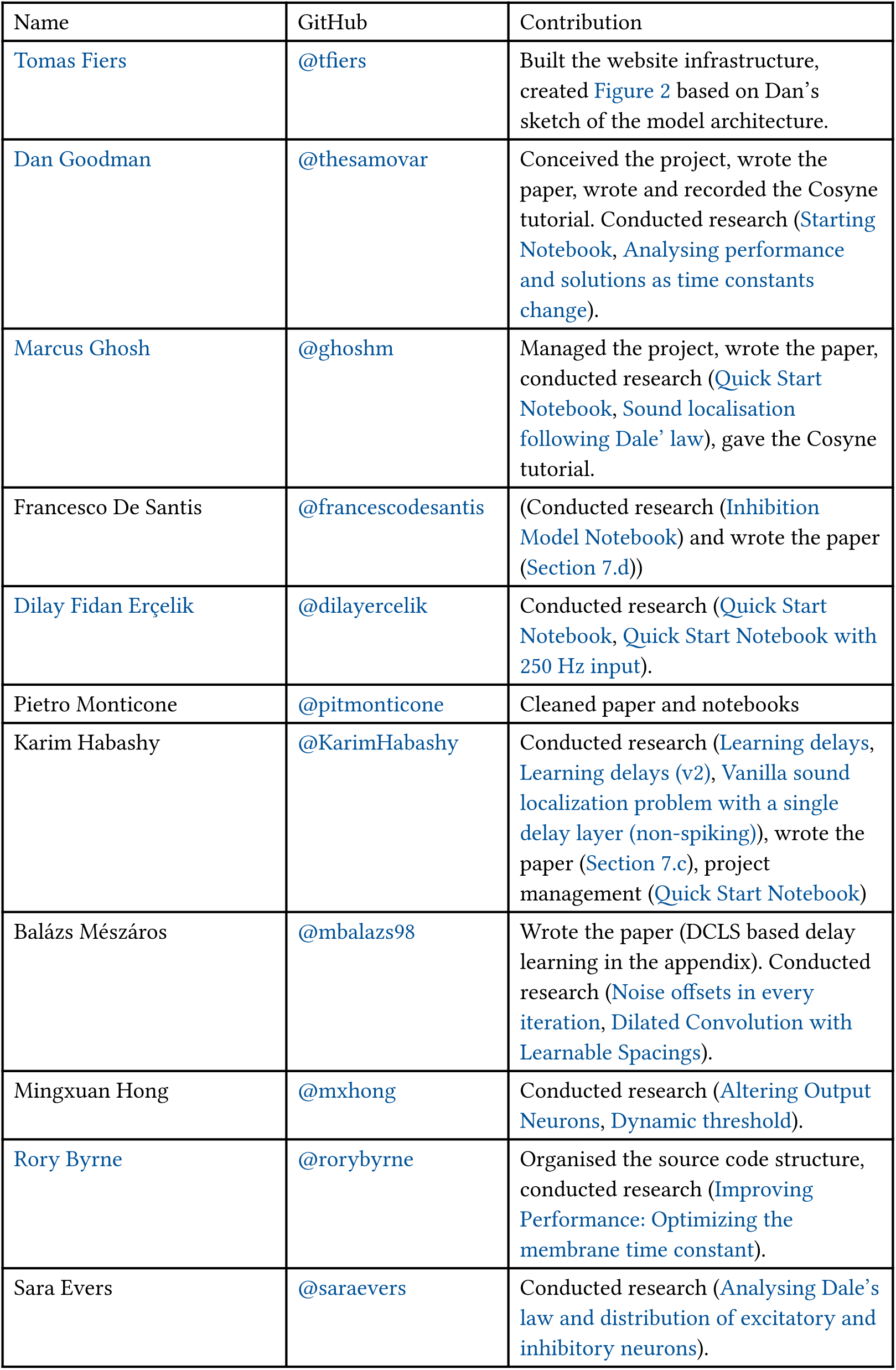
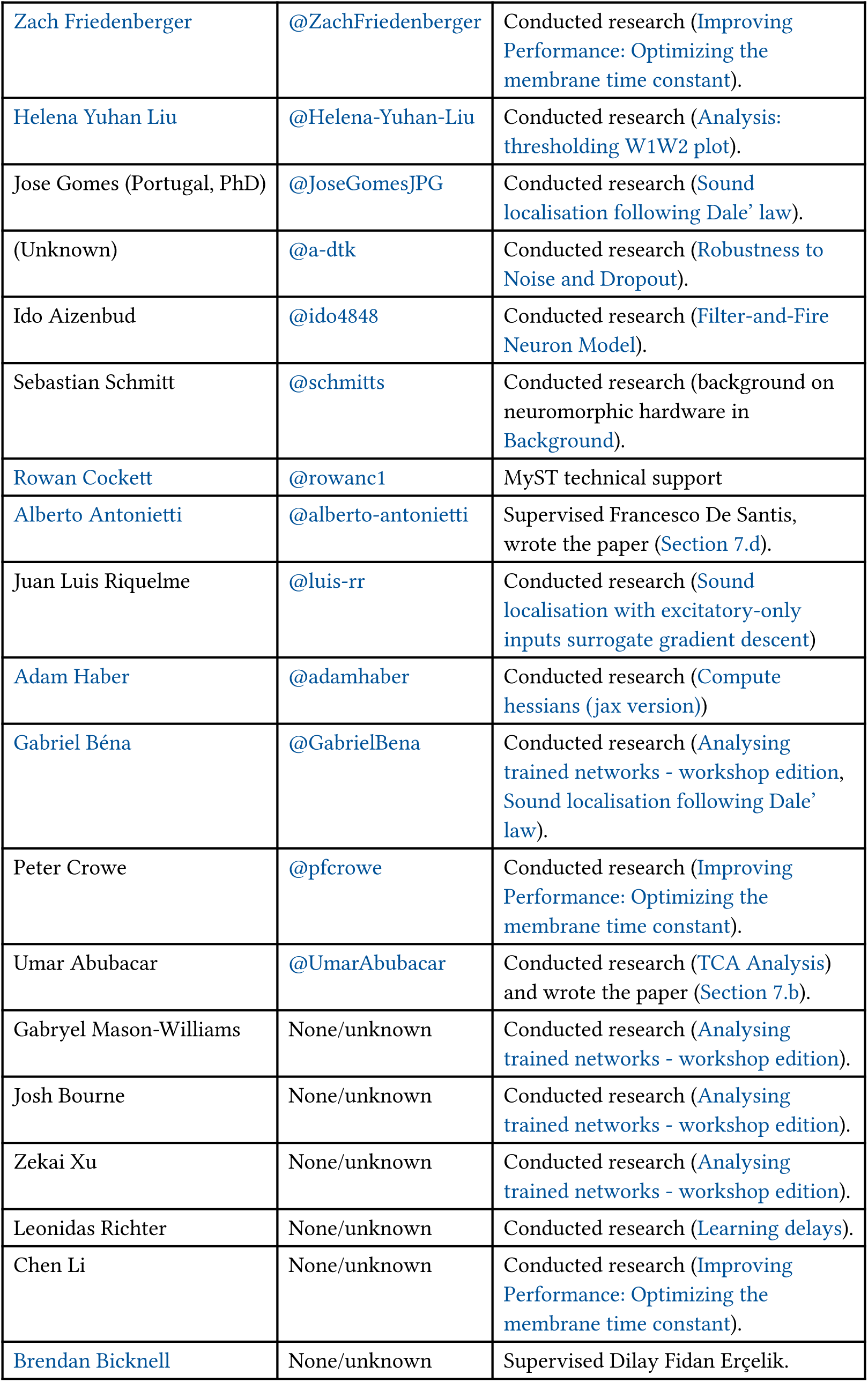
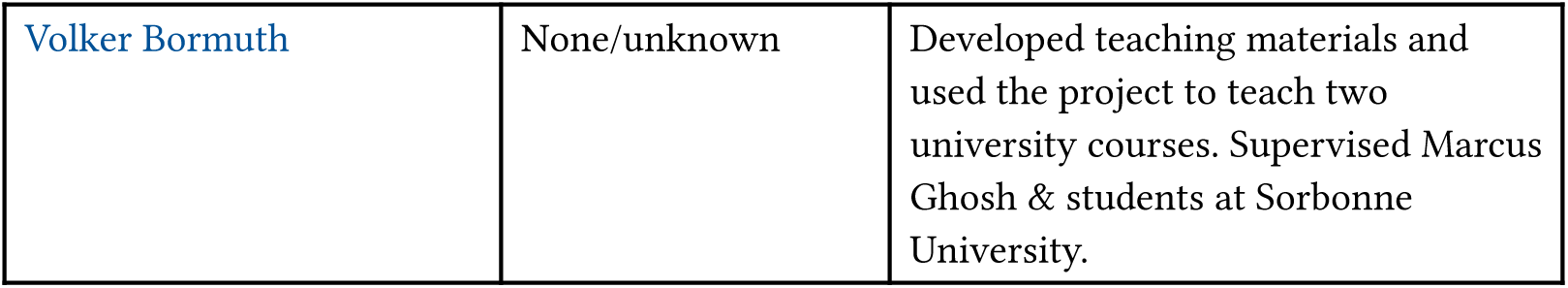
Contributors, ordered by GitHub commits as of 2024-07-16.

| Name | GitHub | Contribution |
| --- | --- | --- |
| Tomas Fiers | @tfiers | Built the website infrastructure, created Figure 2 based on Dan's sketch of the model architecture. |
| Dan Goodman | @thesamovar | Conceived the project, wrote the paper, wrote and recorded the Cosyne tutorial. Conducted research (Starting Notebook, Analysing performance and solutions as time constants change). |
| Marcus Ghosh | @ghoshm | Managed the project, wrote the paper, conducted research (Quick Start Notebook, Sound localisation following Dale' law), gave the Cosyne tutorial. |
| Francesco De Santis | @francescodesantis | (Conducted research (Inhibition Model Notebook) and wrote the paper (Section 7.d)) |
| Dilay Fidan Erçelik | @dilayercelik | Conducted research (Quick Start Notebook, Quick Start Notebook with 250 Hz input). |
| Pietro Monticone | @pitmonticone | Cleaned paper and notebooks |
| Karim Habashy | @KarimHabashy | Conducted research (Learning delays, Learning delays (v2), Vanilla sound localization problem with a single delay layer (non-spiking)), wrote the paper (Section 7.c), project management (Quick Start Notebook) |
| Balázs Mészáros | @mbalazs98 | Wrote the paper (DCLS based delay learning in the appendix). Conducted research (Noise offsets in every iteration, Dilated Convolution with Learnable Spacings). |
| Mingxuan Hong | @mxhong | Conducted research (Altering Output Neurons, Dynamic threshold). |
| Rory Byrne | @rorybyrne | Organised the source code structure, conducted research (Improving Performance: Optimizing the membrane time constant). |
| Sara Evers | @saraevers | Conducted research (Analysing Dale's law and distribution of excitatory and inhibitory neurons). |
| Zach Friedenberger | @ZachFriedenberger | Conducted research (Improving Performance: Optimizing the membrane time constant). |
| Helena Yuhan Liu | @Helena-Yuhan-Liu | Conducted research (Analysis: thresholding W1W2 plot). |
| Jose Gomes (Portugal, PhD) | @JoseGomesJPG | Conducted research (Sound localisation following Dale' law). |
| (Unknown) | @a-dtk | Conducted research (Robustness to Noise and Dropout). |
| Ido Aizenbud | @ido4848 | Conducted research (Filter-and-Fire Neuron Model). |
| Sebastian Schmitt | @schmitts | Conducted research (background on neuromorphic hardware in Background). |
| Rowan Cockett | @rowanc1 | MyST technical support |
| Alberto Antonietti | @alberto-antonietti | Supervised Francesco De Santis, wrote the paper (Section 7.d). |
| Juan Luis Riquelme | @luis-rr | Conducted research (Sound localisation with excitatory-only inputs surrogate gradient descent) |
| Adam Haber | @adamhaber | Conducted research (Compute Hessians (jax version)) |
| Gabriel Béna | @GabrielBena | Conducted research (Analysing trained networks - workshop edition, Sound localisation following Dale' law). |
| Peter Crowe | @pfcrowe | Conducted research (Improving Performance: Optimizing the membrane time constant). |
| Umar Abubacar | @UmarAbubacar | Conducted research (TCA Analysis) and wrote the paper (Section 7.b). |
| Gabryel Mason-Williams | None/unknown | Conducted research (Analysing trained networks - workshop edition). |
| Josh Bourne | None/unknown | Conducted research (Analysing trained networks - workshop edition). |
| Zekai Xu | None/unknown | Conducted research (Analysing trained networks - workshop edition). |
| Leonidas Richter | None/unknown | Conducted research (Learning delays). |
| Chen Li | None/unknown | Conducted research (Improving Performance: Optimizing the membrane time constant). |
| Brendan Bicknell | None/unknown | Supervised Dilay Fidan Erçelik. |
| Volker Bormuth | None/unknown | Developed teaching materials and used the project to teach two university courses. Supervised Marcus Ghosh & students at Sorbonne University. |

## 6. Notebook map

The following lists the notebooks, authors, summary and related notebooks in this project.

### 6.a. Introductory notebooks

**Background** Explanation of the background. (Author: Dan Goodman.)

**Questions & challenges** List of research questions and challenges. (Author: everyone.)

### 6.b. Templates / starting points

**Starting Notebook** The template notebook suggested as a starting point, based on the Cosyne tutorial that kicked off this project. (Author: Dan Goodman.)

**Quick Start Notebook** Condensed version of Starting Notebook using the shorter membrane time constants from Improving Performance: Optimizing the membrane time constant and Dale’s law from Sound localisation following Dale’ law. (Author: Dilay Fidan Erçelik, Karim Habashy, Marcus Ghosh.)

### 6.c. Individual notebooks

**Filter-and-Fire Neuron Model** Using an alternative neuron model. (Author: Ido Aizenbud based on work from Dilay Fidan Erçelik.)

**Altering Output Neurons** Comparison of three different ways of reading out the network’s decision (average membrane potential, maximum membrane potential, spiking outputs) with short and long time constants. (Author: Mingxuan Hong.)

**Analysing trained networks - workshop edition** Group project from an early workshop looking at hidden unit spiking activity and single unit ablations. Found that some hidden neurons don’t spike, and ablating those does not harm performance. Builds on (WIP) Analysing trained networks. (Author: Gabriel Béna, Josh Bourne, Tomas Fiers, Tanushri Kabra, Zekai Xu.)

**Sound localisation following Dale’ law** Investigation into the results of imposing Dale’s law. Incorporated into Quick Start Notebook. Uses a fix from Analysing Dale’s law and distribution of excitatory and inhibitory neurons. (Author: Marcus Ghosh, Gabriel Béna, Jose Gomes.)

**Dynamic threshold** Adds an adaptive threshold to the neuron model and compares results. Conclusion is that the dynamic threshold does not help in this case. (Author: Mingxuan Hong.)

**Sound localisation with excitatory-only inputs surrogate gradient descent** Results of imposing an excitatory only constraint on the neurons. Appears to find solutions that are more like what would be expected from the Jeffress model. (Author: Juan Luis Riquelme.)

**Learning delays, Learning delays (v2) and Vanilla sound localization problem with a single delay layer (non-spiking)** Delay learning using differentiable delay layer, written up in Section 7.c (Author: Karim Habashy.)

**Dilated Convolution with Learnable Spacings** Delay learning using Dilated Convolution with Learnable Spacings, written up in Section 7.c. (Author: Balázs Mészáros.)

**Robustness to Noise and Dropout** Test effects of adding Gaussian noise and/or dropout during training phase. Conclusion is that dropout does not help and adding noise decreases performance. (Author: Unknown (@a-dtk).)

**Version with 250 Hz input, Quick Start Notebook with 250 Hz input** Analysis of results with a higher frequency input stimulus and different membrane time constants for hidden and output layers. Conclusion is that smaller time constant matters for hidden layer but not for output layer. (Author: Dilay Fidan Erçelik.)

**Analysing performance and solutions as time constants change** Deeper analysis of strategies found by trained networks as time constants vary. Added firing rate regularisation. Extends Improving Performance: Optimizing the membrane time constant. Written up in more detail in Section 7.a. (Author: Dan Goodman.)

**Workshop 1 Write-up** Write-up of what happened at the first workshop. (Author: Marcus Ghosh.)

### 6.d. Inconclusive

The following notebooks did not reach a solid conclusion.

**Compute hessians (jax version)** An unfinished attempt to perform sensitivity analysis using Hessian matrices computed via autodifferentiation with the Jax library. (Author: Adam Haber.)

**Noise offsets in every iteration** Analysis of an alternative way of handling noise. (Author: Balázs Mészáros.)

**Analysis: thresholding W1W2 plot** Unfinished attempt to improve analysis code. (Author: Helena Yuhan Liu.)

### 6.e. Historical

This subsection includes notebooks whose content got merged into an updated notebook later.

**(WIP) Analysing trained networks** Early work on analysing the strategies learned by trained networks. Folded into Analysing trained networks - workshop edition. (Author: Dan Goodman.)

**Improving Performance: Optimizing the membrane time constant** Analyses how performance depends on membrane time constant. Folded into Analysing performance and solutions as time constants change. (Author: Zach Friedenberger, Chen Li, Peter Crowe.)

**Analysing Dale’s law and distribution of excitatory and inhibitory neurons** Fixed a mistake in an earlier version of Sound localisation following Dale’ law. (Author: Sara Evers.)

## 7. Appendices

In this section, we provide detailed write-ups of our scientific results.

### 7.a. A minimal trainable model of IPD processing

|  |  |
| --- | --- |
| Section authors | Dan Goodman |
| Notebooks | Analysing performance and solutions as time constants change |

This section describes the initial model developed for the Cosyne tutorial that served as the starting point for this project (Goodman et al., 2022). It also describes some small variants of this basic model produced in the course of the project, which can be seen in the notebook Analysing performance and solutions as time constants change.

The aim of the model was to address a long standing question about how and why the brain localises sounds in the way it does, restricted specifically in this case to interaural phase differences (IPDs) cues used at low frequencies. A common strand in this research is to consider a population of binaurally responsive neurons, each with a different frequency and IPD tuning, summarised by their best frequency (BF) and best delay (BD), i.e. the frequency and delay/phase at which they give their strongest response. From this spatial-temporal *encoding* of the sound, a second network attempts to *decode* the sound location. In the place theory of Jeffress (1948), the encoding is done by coincidence detection neurons arrayed so that each neuron receives a sound from the left and right ear with different conduction delays. Decoding proceeds as follows. When the conduction delays match the acoustic delays induced by the arrival time difference of the sound at the two ears, the neuron fires at a maximal rate because of the coincidence detection. Doubt was cast on this theory by McAlpine et al. (2001), who argued that it was not robust to neural noise, and proposed instead a “hemispheric code” that encodes the sound in the difference in the average firing rates of neurons whose best delay is positive versus negative. While this optimises robustness to neural noise, Goodman et al. (2013) showed that it was not efficient at integrating across frequencies, was biased in the presence of acoustic noise, and generalised poorly to sounds outside of the training set.

In this project, we asked what strategies would be found when jointly optimising for both encoding and decoding using modern machine learning methods to train a network end-to-end.

#### 7.a.i. Methods

The model consists of the following pathway, illustrated in Figure 3: IPD → stimulus → input neurons → hidden layer neurons → readout neurons.

**Figure 3:**
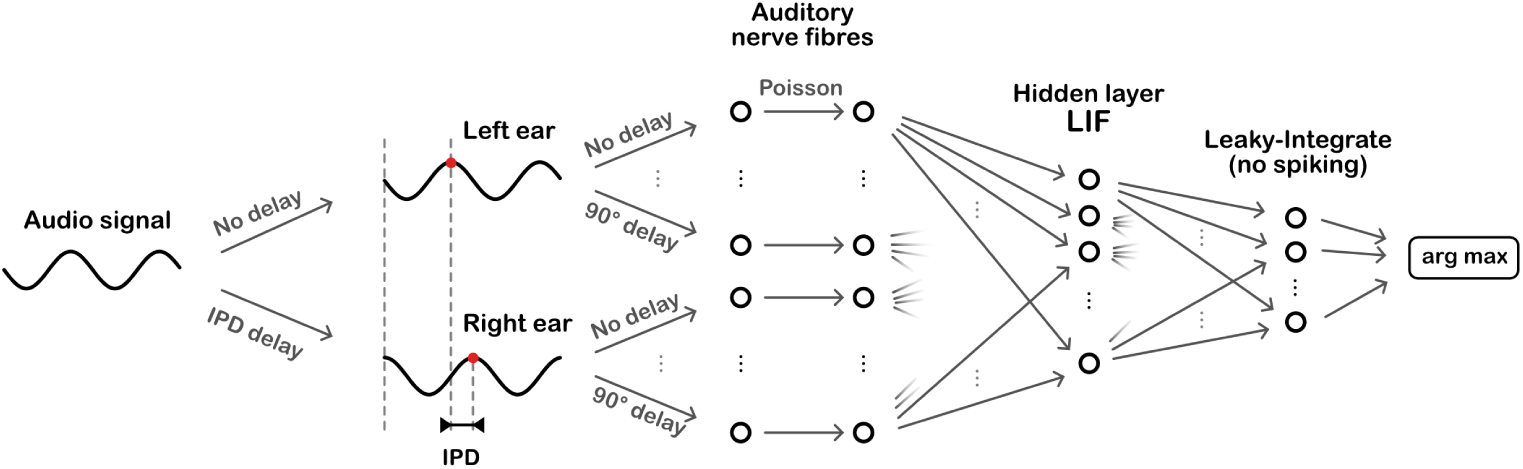
Overall model architecture. Sinusoidal audio signals are passed through two populations of units, with a range of preassigned delays, representing the left and right ears. These units generate Poisson spike trains which pass forward to a layer of leaky-integrate and fire (LIF) units, then a layer of leaky-integrator output units from which we readout the networks estimate of the interaural phase difference (IPD).

The IPD is an angle uniformly randomly selected in ɑ ∈ [−π/2, π/2] (frontal plane only).

The stimulus is a sine wave, and we model interaural phase differences (IPDs) by adding the IPD ɑ as a phase delay to the right ear. Enumerating the ears with index 𝑖 ∈ {0, 1} so that 𝑖 = 0 is the left ear, we get that the stimulus in ear 𝑖 is:

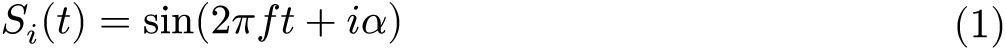

To model a population of spiking neurons with different lags, we associate each input layer neuron with an ear and an additional phase delay 𝜓 uniformly spaced in the range (0, π/2). This ensures that by comparing a left and right ear you can generate any phase difference in the range (−π/2, π/2) as required. Specifically, we set neuron *j* connected to ear 𝑖 to receive the stimulus:

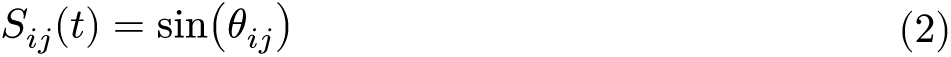

where

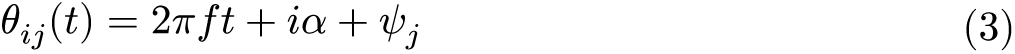

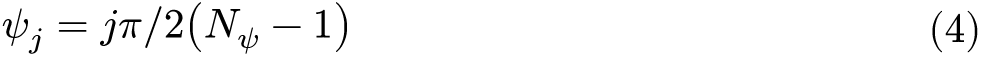

and *N*_𝜓_ is the number of input neurons per ear.

Next, we make these neurons spiking by giving them a time varying firing rate:

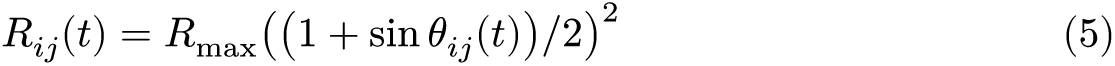

where *R*_max_ is the maximum firing rate. Spikes are then generated via an inhomogeneous Poisson process with intensity *R*_*ij*_(*t*). Some example raster plots are shown in Figure 4.

**Figure 4:**
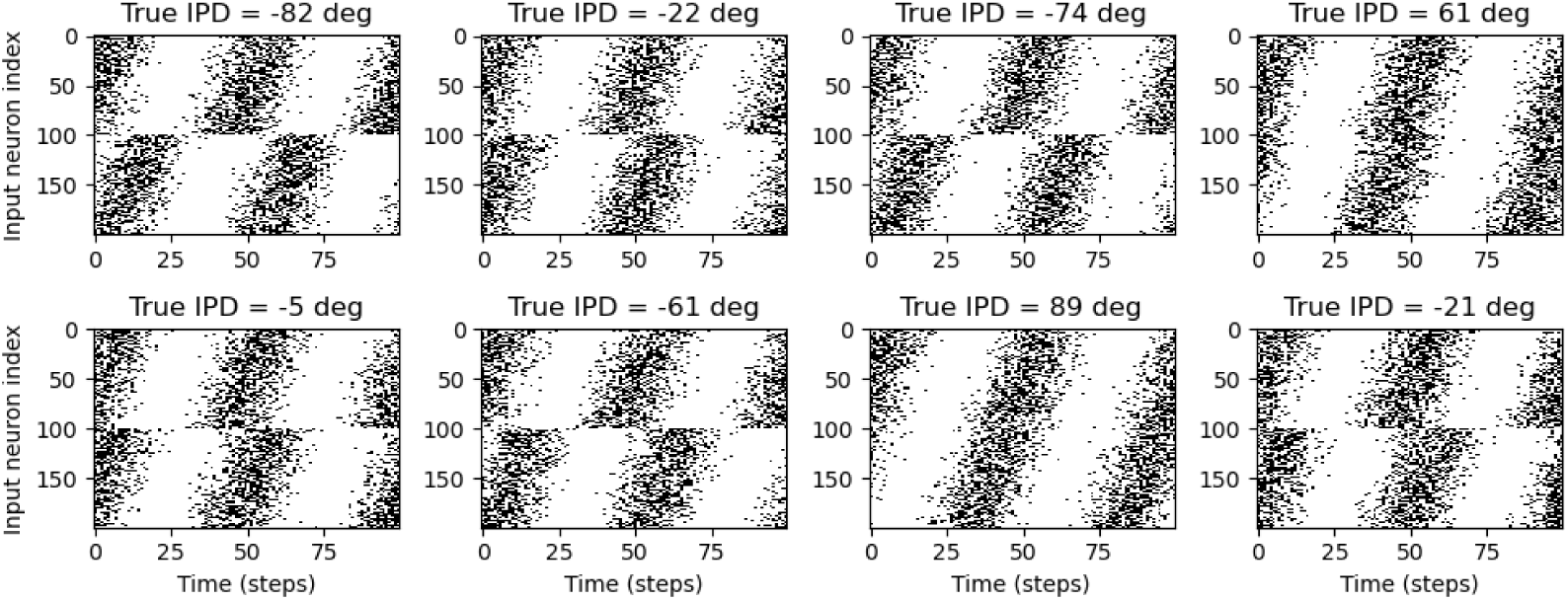
Examples of generated input spike trains.

The 2*N*_𝜓_ input neurons are connected all-to-all to a “hidden layer” of *N*_ℎ_ spiking neurons. These are standard leaky integrate-and-fire neurons with a membrane potential 𝑣 that in the absence of spikes evolves over time according to the differential equation:

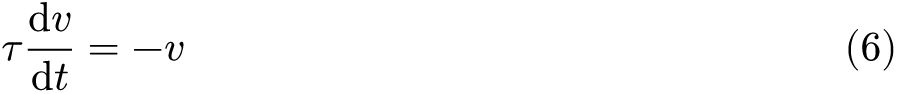

where 𝜏 is the membrane time constant. An incoming spike on a synapse with weight 𝑤 causes an instantaneous increase 𝑣 ← 𝑣 + 𝑤. These weights are stored in a matrix *W*_𝑖ℎ_ of size (2*N*_𝜓_, *N*_ℎ_). If 𝑣 crosses the spike threshold of 1, the neuron emits a spike and instantaneously resets to 𝑣 ← 0.

The hidden layer is all-to-all connected to a readout layer of *N*_𝑐_ neurons via a weight matrix *W*_ℎ𝑜_. The aim is to divide the set of possible IPDs into *N*_𝑐_ intervals *I*_𝑘_ = [−π/2 + 𝑘π/*N*_𝑐_, −π/2 + (𝑘 + 1)π/*N*_𝑐_] and then, if neuron 𝑘 is the most active, guess that the IPD must be in interval *I*_𝑘_. These hidden layer neurons follow the same differential equation but do not spike. Instead, to guess the IPD we compute their mean membrane potential over the duration of the stimulus, | 𝑣_𝑘_, and then compute an output vector that is the log softmax function of these mean membrane potentials:

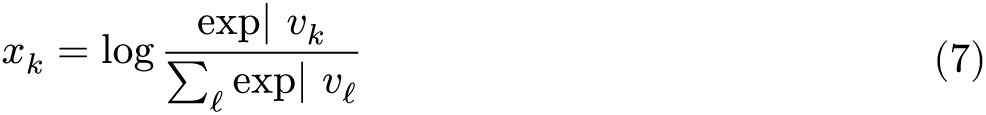

We then interpret *x*_𝑘_ as the estimated probability that ɑ ∈ *I*_𝑘_. Our estimate of the IPD 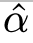 will be the midpoint of the interval corresponding to the most active neuron 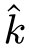 = argmax_𝑘_ *x*_𝑘_. Note that the softmax function and probability interpretation are important for training the network, but once the network is trained you can equally well compute 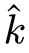 = argmax_𝑘_| 𝑣_𝑘_.

The network is trained by defining a loss function that increases the further away the network behaviour is from what we would like (defined in detail below), and then using the surrogate gradient descent method (Neftci et al., 2019; Zenke & Ganguli, 2018). Full details on training parameters can be found in the notebook Starting Notebook.

The loss function we use is composed of two terms. The first is the cross entropy or negative log likelihood loss that measures how far our predicted probability distribution *x*_𝑘_ is from the true probability distribution (which has value 1 for the correct 𝑘 and 0 for all other 𝑘). The second term, which is not used in all the notebooks in this project, is an optional regularisation term. In Analysing performance and solutions as time constants change we regularise based on the firing rates of the hidden layer neurons. We compute the firing rate for each hidden neuron *r*_*m*_. If this is below a mimimum threshold *r*_−_ it contributes nothing to the loss, otherwise we compute *L*_*m*_ = ((*r*_*m*_ − *r*_−_)/(*r*_+_ − *r*_−_))^2^ for each neuron for a constant *r*_+_ explained below. We now compute the average and multiply a constant *L* = 𝑐 ∑_*m*_ *L*_*m*_/*N*_ℎ_. The constant *r*_+_ is the maximum firing rate we would like to see in the network, so that *L*_*m*_ = 1 if *r*_*m*_ = *r*_+_. The constant 𝑐 is chosen to be the expected initial cross-entropy loss of the network before training. This makes sure that a firing rate of *r*_*m*_ = *r*_+_ is heavily penalised relative to the cross-entropy loss, but that any firing rate below *r*_−_ is fine. We chose *r*_−_ = 100 sp/s and *r*_+_ = 200 sp/s.

#### 7.a.ii. Results

This approach is able to train a network that can perform the task well, using very few neurons (*N*_𝜓_ = 100 input neurons per ear, *N*_ℎ_ = 8 hidden neurons and *N*_𝑐_ = 12 output neurons). Mean absolute error in IPD is ∼ 2.6 deg (Figure 5). Hidden neuron firing rates are between 110 and 150 sp/s (Figure 6).

**Figure 5:**
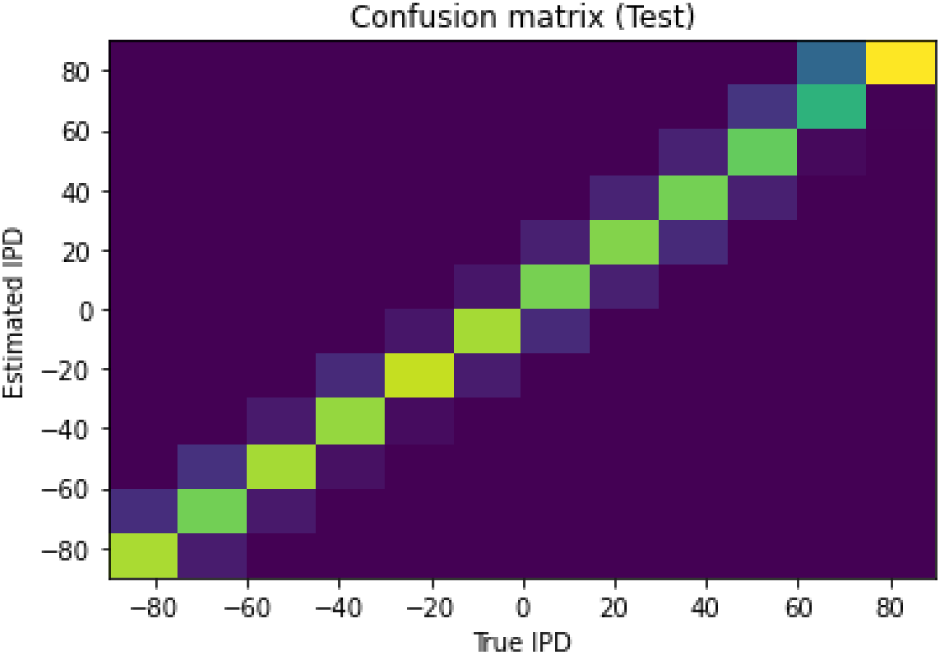
Confusion matrix. Results of training the network with 𝑓 = 50 Hz, 𝜏 = 2 ms, *N*_𝜓_ = 100, *N*_ℎ_ = 8, *N*_𝑐_ = 12. Mean absolute IPD errors are ∼ 2.6 deg.

**Figure 6:**
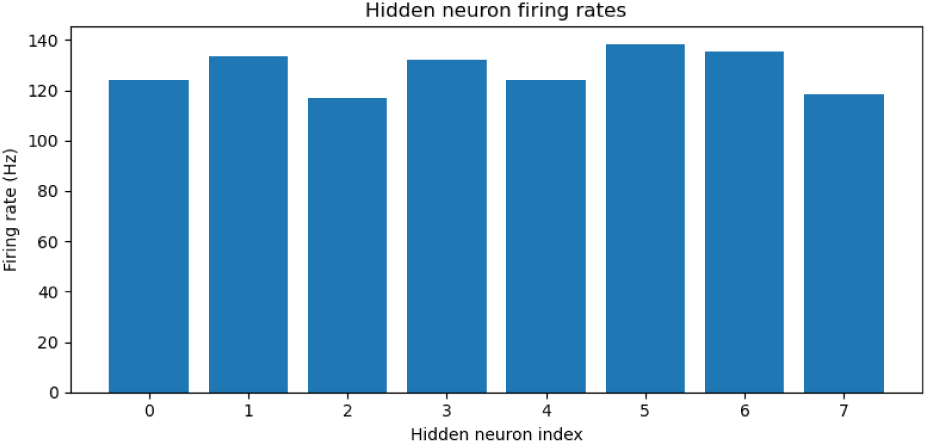
Hidden neuron firing rates. Results of training the network with 𝑓 = 50 Hz, 𝜏 = 2 ms, *N*_𝜓_ = 100, *N*_ℎ_ = 8, *N*_𝑐_ = 12. Mean absolute IPD errors are ∼ 2.6 deg.

Analysis of the trained networks show that it uses an unexpected strategy. Firstly, the hidden layer neurons might have been expected to behave like the encoded neurons in Jeffress’ place theory, and like recordings of neurons in the auditory system, with a low baseline response and an increase for a preferred phase difference (best phase). However, very reliably they find an inverse strategy of having a high baseline response with a reduced response at a least preferred phase difference (Figure 7). Note that the hidden layer neurons have been reordered in order of their least preferred delay to highlight this structure. These shapes are consistently learned, but the ordering is random. By contrast, the output neurons have the expected shape (Figure 8). Interestingly, the tuning curves are much flatter at the extremes close to an IPD of ±π/2. We can get further insight into the strategy found by plotting the weight matrices *W*_𝑖ℎ_ from input to hidden layer, and *W*_ℎ𝑜_ from hidden layer to output, as well as the product *W*_𝑖𝑜_ = *W*_𝑖ℎ_ ⋅ *W*_ℎ𝑜_ which would give the input-output matrix for a linearised version of the network (Figure 9).

**Figure 7:**
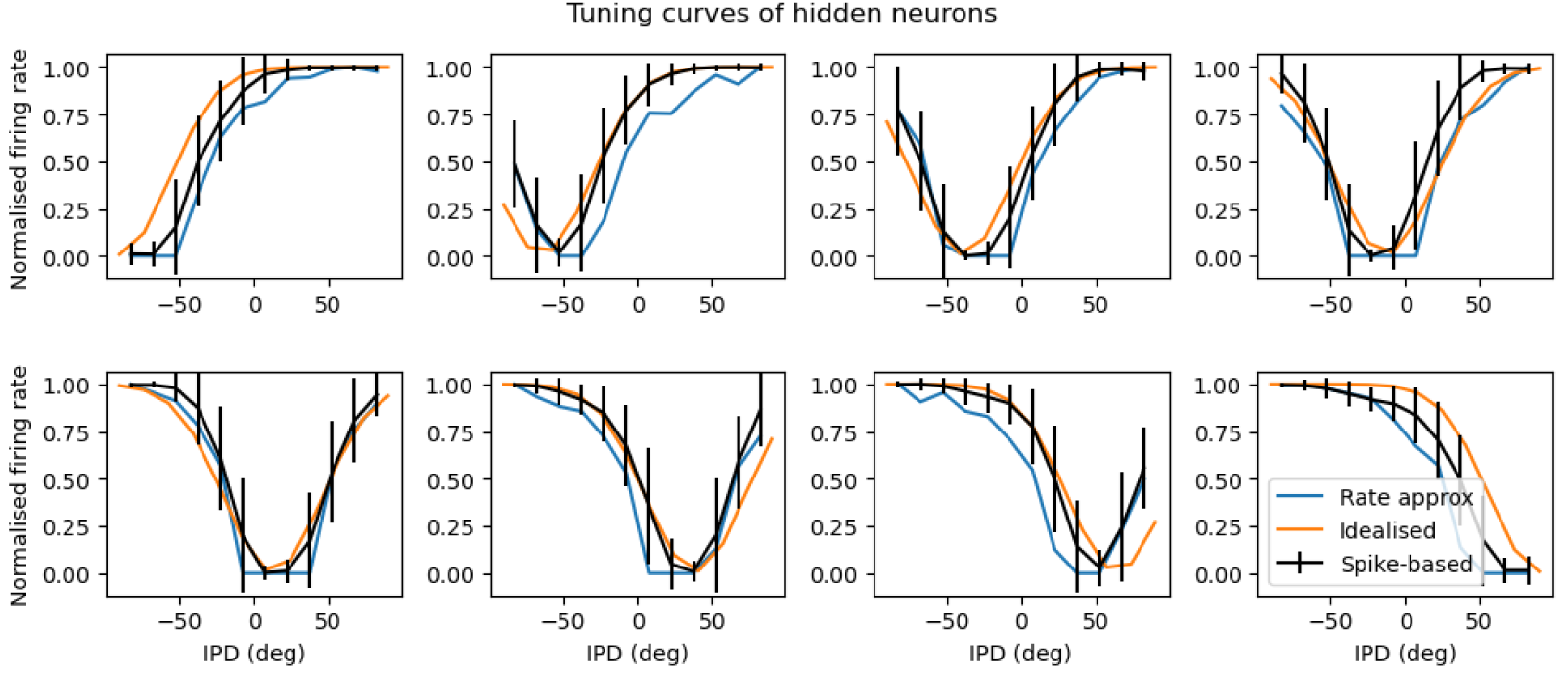
Tuning curves of hidden neurons. Strategy found by trained network with 𝑓 = 50 Hz, 𝜏 = 2 ms, *N*_𝜓_ = 100, *N*_ℎ_ = 8, *N*_𝑐_ = 12. *N*_𝜓_ = 100, *N*_ℎ_ = 8, *N*_𝑐_ = 12.

**Figure 8:**
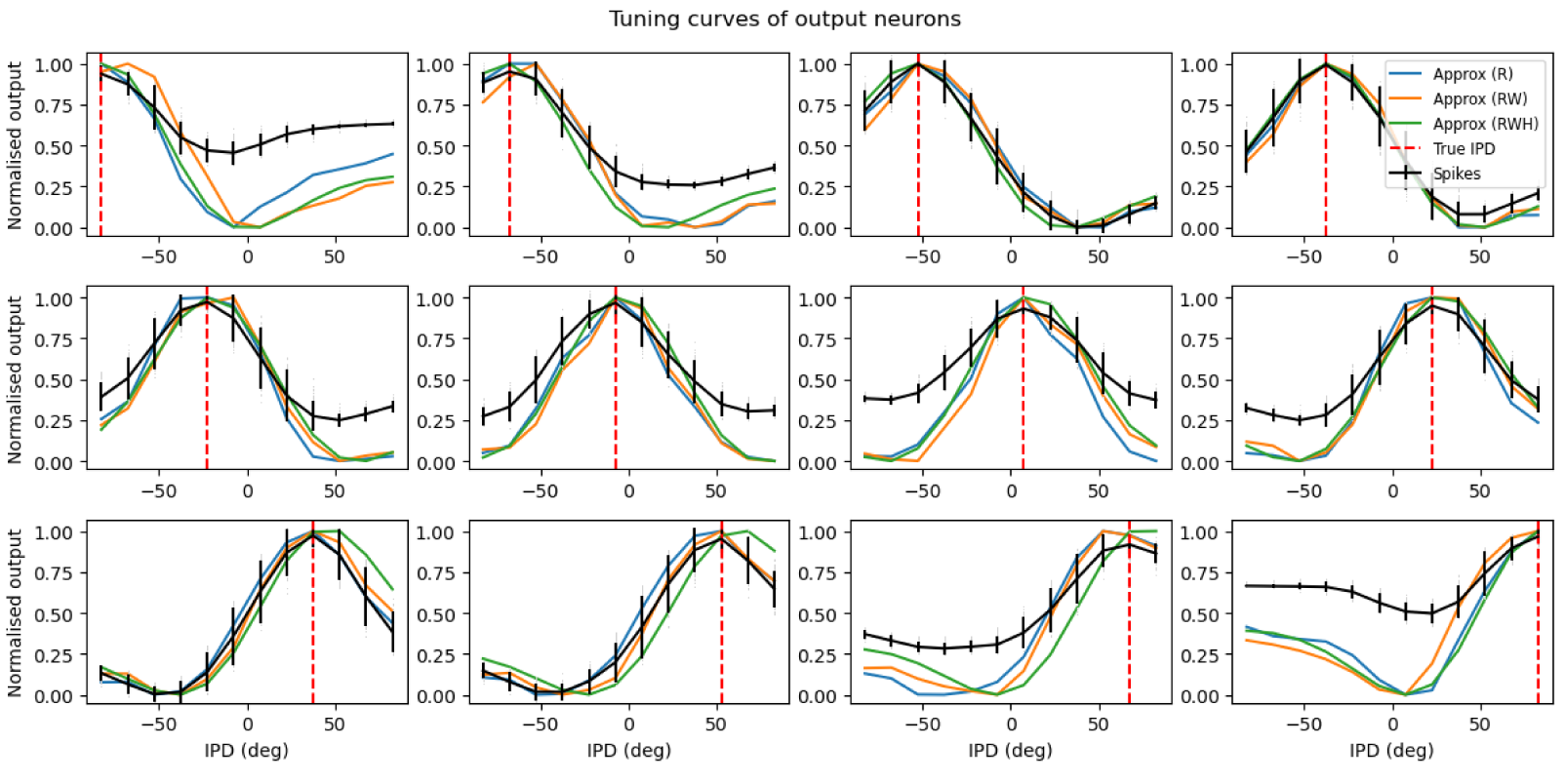
Tuning curves of hidden neurons. The dashed red lines indicate the estimated IPD if that neuron is the most active.Strategy found by trained network with 𝑓 = 50 Hz, 𝜏 = 2 ms,

**Figure 9:**
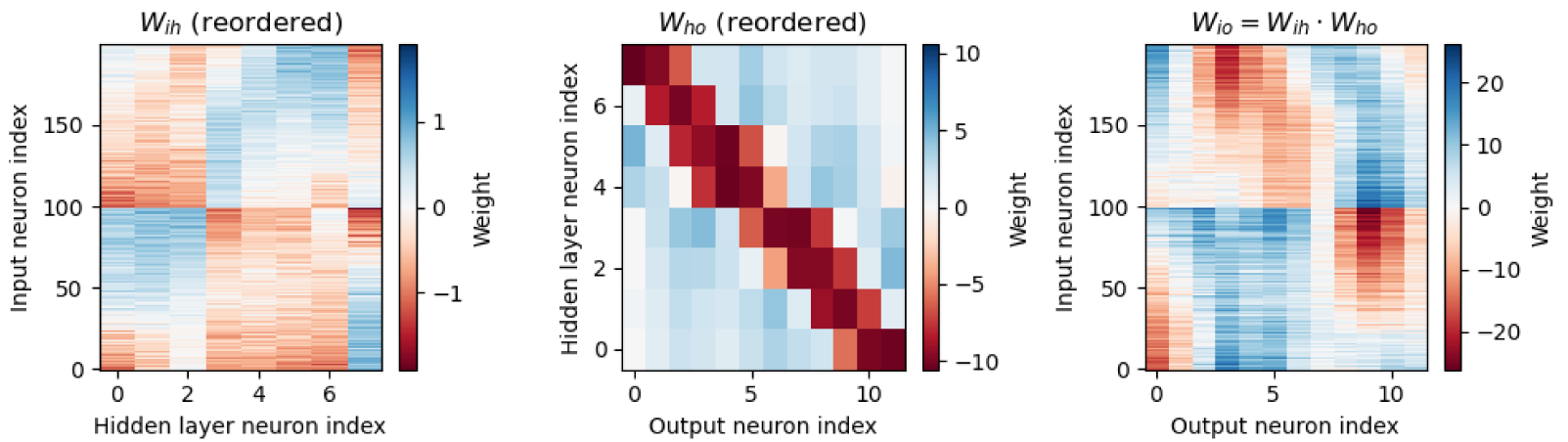
Weight matrices, with hidden neurons reordered by best delay. Strategy found by trained network with 𝑓 = 50 Hz, 𝜏 = 2 ms, *N*_𝜓_ = 100, *N*_ℎ_ = 8, *N*_𝑐_ = 12.

A number of features emerge from this analysis. The first is that the tuning curves of the hidden neurons have a very regular structure of having a high baseline firing rate with a dip around a “least preferred” delay that varies uniformly in the range −π/2 to π/2. Indeed, the tuning curves 𝑖 can be very well fit with the function 𝑎 + 𝑏𝑒^−(ɑ−ɑ𝑖)2/2𝜎2^ where ɑ is the IPD, ɑ_𝑖_ = −π/2 + 𝑖π/*N*_ℎ_ is the “least preferred” IPD, and 𝑎, 𝑏, 𝜎_ɑ_ are parameters to fit (Figure 7, orange lines). This would look likely to be consistent with some form of optimal coding theory that minimises the effect of the Poisson noise in the spike counts, although we did not pursue this explanation.

The second feature is that spike timing does not appear to play a significant role in this network. This may initially appear suprising but in fact it is inevitable because of the coding scheme we have used where the initial layer of neurons fire Poisson spikes, and there is only one spiking layer, meaning there is no scope for spike times to be used (a limitation of this model realised late in the process). Indeed, if we predict the output of the network purely using the firing rates of the input stimulus passed through the weight matrices *W*_𝑖ℎ_ and *W*_ℎ𝑜_ plus a static nonlinearity for the input to hidden layer, we get an excellent approximation for the hidden neurons (Figure 7, blue lines) and a qualitatively good fit for the output neurons (Figure 8, blue lines). Specifically, if *r*_𝑖_(*t*) is the instantaneous time-varying firing rate of input neuron 𝑖 we approximate the instantaneous hidden units firing rates by the function:

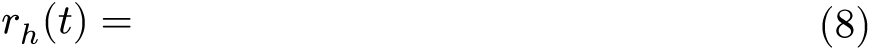

This function is the firing rate of a leaky integrate-and-fire neuron 𝜏 𝑣^′^ = *r*_𝑖_(*t*) − 𝑣 with a refractory period *t*_refrac_ (which is d*t* in our case because of the way it is simulated) if the function *r*_𝑖_(*t*) were constant over time, but it fits well even with a time-varying *r*_𝑖_(*t*). From this we can take an average over time to get the mean firing rate. The output units do not spike so their activity is simply approximated by *r*_𝑜_(*t*) = ∑_ℎ_ *W*_ℎ𝑜_*r*_ℎ_(*t*).

Finally, we note that the weight matrix *W*_ℎ𝑜_ visible in Figure 9 seems to have a very regular structure of weights that have a broad excitation and a narrowly tuned inhibition (an unusual pattern). Indeed, we can fit this well with a Ricker wavelet (or “Mexican hat”) function:

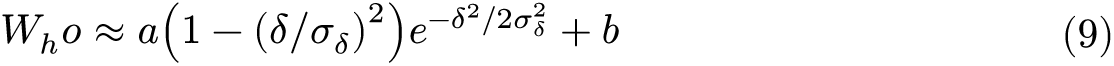

where 𝛿 = 𝑜 − *N*_𝑐_ℎ/*N*_ℎ_, ℎ and 𝑜 are the indices of the hidden and output neurons, *N*_ℎ_ is the number of hidden neurons, *N*_𝑐_ the number of output neurons, and 𝑎, 𝑏 and 𝜎_𝛿_ are parameters to estimate. Using this approximation and the rate-based approximation from before, we get the orange curves in Figure 8. If we use both the Ricker wavelet approximation of *W*_ℎ𝑜_ and the idealised tuning curves, we get the green curves. All in all, this gives us a 6 parameter model that fits the data extremely well, a significant reduction on the 896 parameters for the full model (*N*_𝜓_*N*_ℎ_ + *N*_ℎ_*N*_𝑐_).

#### 7.a.iii. Discussion

This subproject was an extension of the original notebook Starting Notebook with the aim of understanding the solutions found in more detail. We successfully found a 6-parameter reduced model that behaves extremely similarly to the full model, and we can therefore say that we have largely understood the nature of this solution. We did not look in detail for a deep mathematical reason why this is the solution that is found, and this would make for an interesting follow-up. Are these tuning curves and weights Bayes optimal to reduce the effect of the Poisson spiking noise, for example?

The solution that was found gives tuning curves that are unlike those found in natural auditory systems, with an inverted “least preferred phase” structure instead of the typical “preferred phase”. In addition, the weight matrix from the hidden to output layer has broadly tuned excitation and narrowly tuned inhibition, which is an unusual pattern.

The solution found does not appear to use coincidence detection properties of spiking neurons, and indeed can be well approximated by a purely rate-based approximation. It appears to find a solution similar to that suggested by the equalisation-cancellation theory (Culling, 2020; Durlach, 1963). This seems initially surprising, but in fact because of the nature of the input stimulus (Poisson spike trains) and the fact that there is only one spiking layer of neurons, there is no temporal structure for coincidence detection to make use of, so it was inevitable that it would not find a solution that uses this strategy. An interesting follow-up would be to use a more detailed model of neuronal firing in the cochlear nucleus for example, or a multi-layer structure, and see if different solutions are found.

### 7.b. Tensor component analysis

|  |  |
| --- | --- |
| Section authors | Umar Abubacar |
| Notebooks | TCA Analysis |

We used tensor component analysis (TCA) to explore the representation and learning dynamics of multiple models by examining their neuron, trial, and temporal factors. TCA helps decompose multidimensional data into core components that capture the essential structure and relationships inherent in the neural activities across different dimensions. This analysis provides a clear insight into how models encode and process information over multiple trials, revealing the underlying mechanisms that drive their performance and behaviour. By employing hierarchical clustering with Ward’s method, we focused on assessing the similarity in neuron factor representations across models, which has elucidated the consistency and diversity of neural encoding across different training instances.

#### 7.b.i. Results

We use nonnegative tensor component analysis (TCA) to examine spiking activity from the hidden layer of a neural network. Gaussian smoothing is applied to the spike events, and trials are stacked into a single tensor. The rank-1 decomposition in Figure 10 during training illustrates the learning process of the model, characterised by increased activity across all IPD bins through a subset of hidden neurons. The analysis highlights predominant firing within two critical intervals in the time domain: 0-25ms and 50-75ms, which aligns with periods of active input signals.

**Figure 10:**
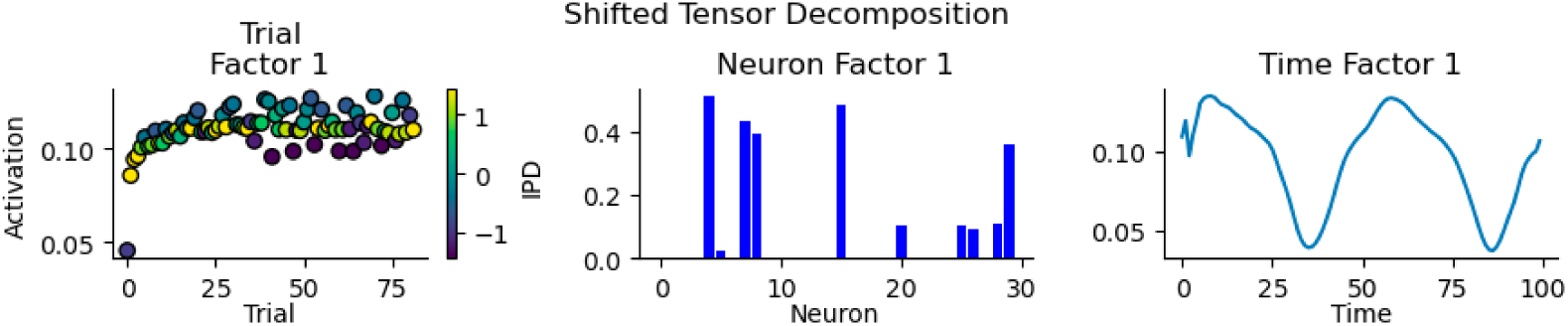
Rank 1 decomposition of spiking during training of the simple model.

To capture the categorical dynamics within the model, the optimal number of components was determined by minimising reconstruction error and maximising component similarity. An optimal count of six ranks was identified for the model and are shown in Figure 11. The factors derived from tensor component analysis reveal distinct patterns of neural activation across time, highlighting the dynamic nature of responses to stimuli. Specifically when looking at trail factors, Factor 5 is responsive to the extreme IPDs of π/2 and −π/2, while Factors 1 and 4 focus on more central IPDs. Factors 2 and 3 indicate neuron assemblies responsive to IPDs towards −π/2 as is Factor 6 to π/2.

**Figure 11:**
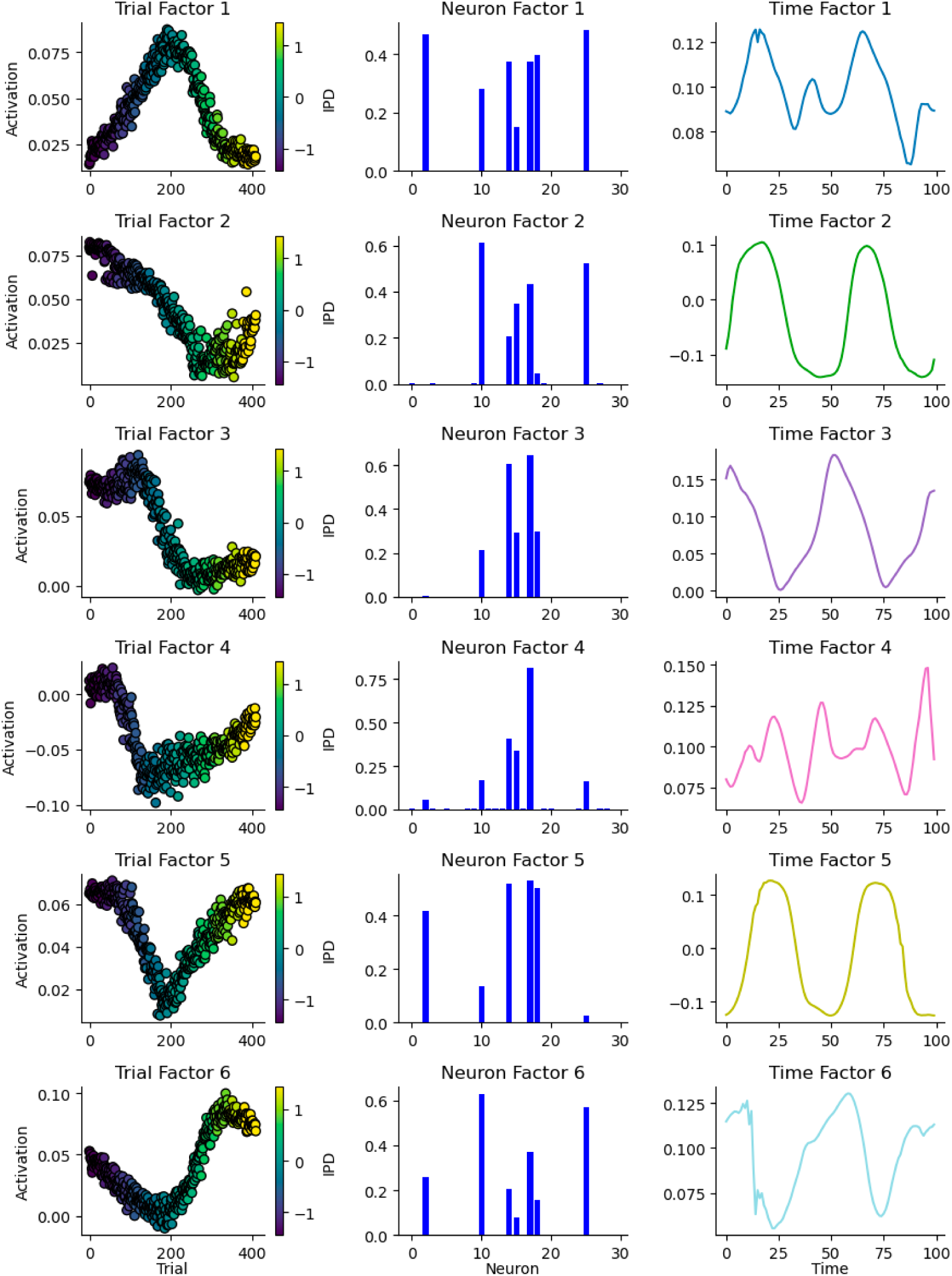
Rank 6 decomposition of spiking during training of the simple model.

#### 7.b.ii. Model variability

To test variability between different instances of the simple model we trained 50 models on the same training data and compared the neuron and temporal factors of the rank 1 decomposition. Neuron factors were sorted by activation strength to align the most significant neurons across models and factors were normalised. Performing clustering analysis of the neuron factors between models helps to elucidate the ensembles responsible for learning the task. The dendrogram from the clustering provided a visual representation of these relationships, with models grouped together showcasing more similar model dynamics compared to those further apart. Here we can see grouping of models into 2 major categories in Figure 12.

**Figure 12:**
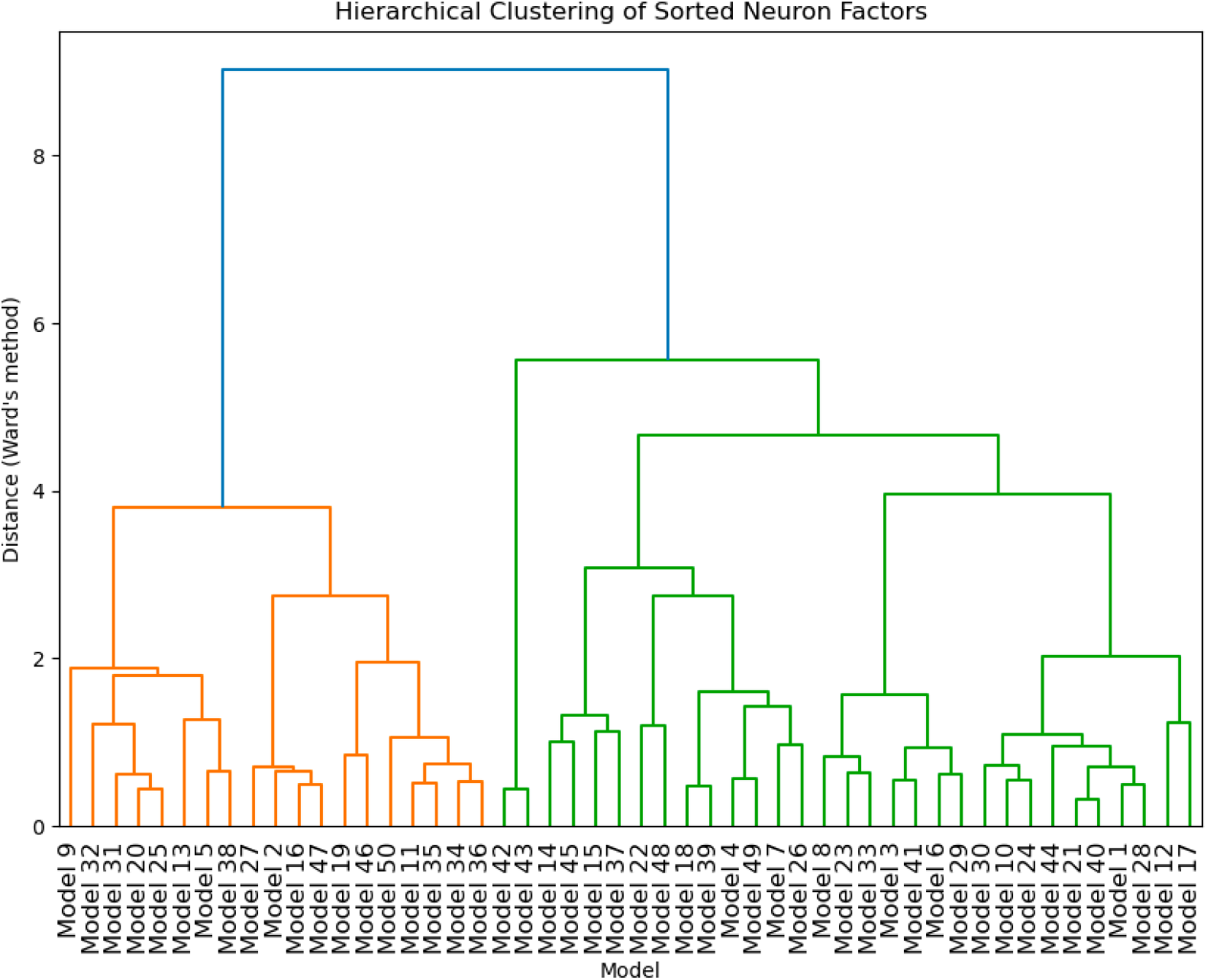
Hierarchical clustering of neuron factors across models.

Sampling models from each cluster shows that clusters with a higher number of active neurons tend to have better training accuracy but do not consistently achieve better test accuracy. In fact, clusters with fewer active neurons sometimes demonstrate better generalisation which had a slightly higher test accuracy despite fewer active neurons. This suggests that models with a moderate number of active neurons can achieve a balance between training effectiveness and test generalisation, potentially leading to more robust solutions. This discrepency in accuracies can be seen in Figure 13 where cluster 1-4 are increase in average active neurons.

**Figure 13:**
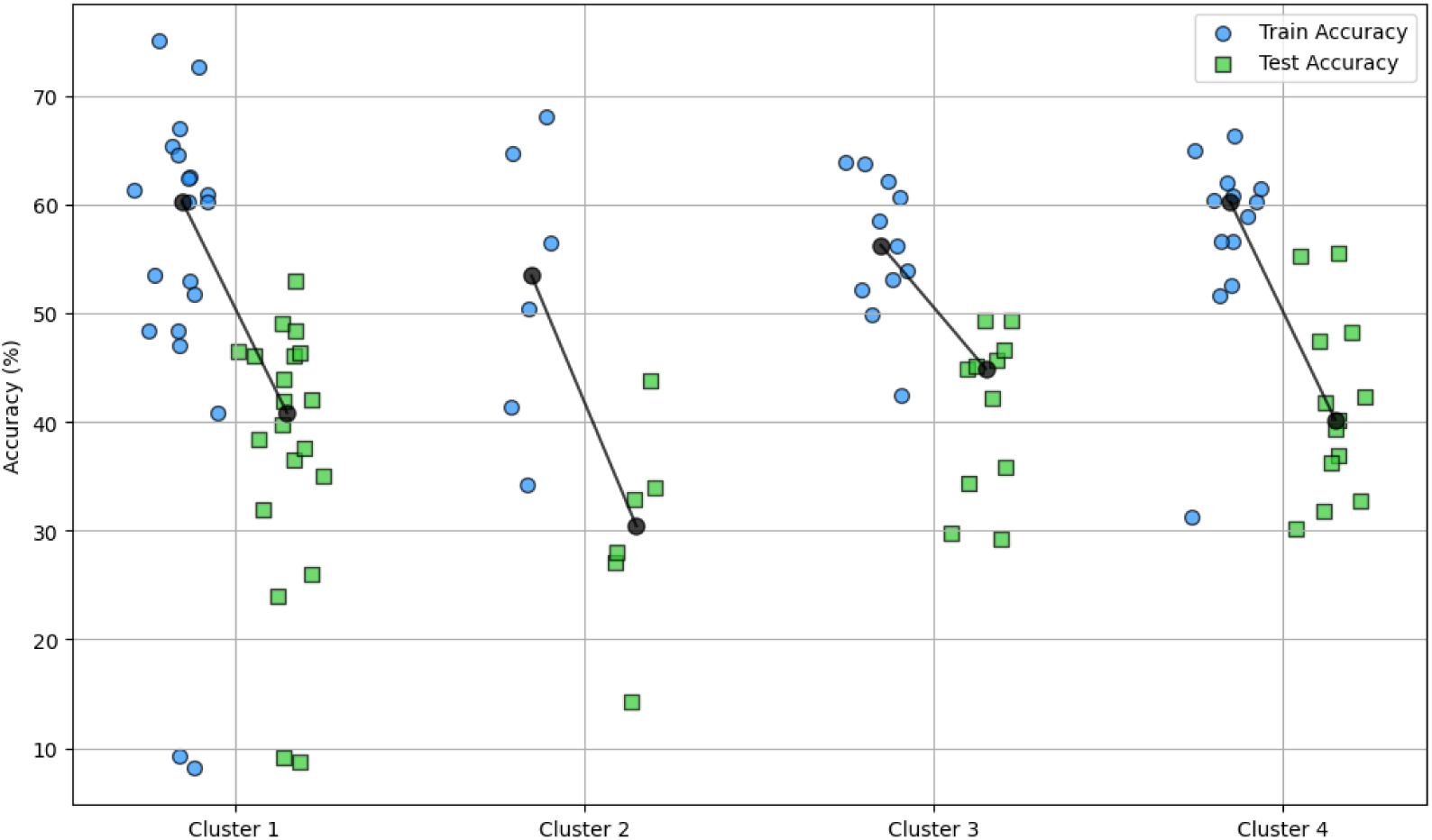
Training and test accuracy of models in each cluster.

Comparing the activation of neural assemblies with more ranks as in Figure 12 shows similar tuning across trails and models suggesting the models learn the same task but are able to do so with a range of subsets of neurons at the cost of accuracy.

Visualising the mean and standard deviation of temporal factors across models shows a common pattern of activation within the hidden layer neurons. The representation in Figure 14 highlights periods of peak neuronal activity. The shifts in peak timings across different models suggest variability in how these models prioritise processing delays related to inputs. Such differences likely stem from the weights learned by each model, reflecting the phase differences prioritised.

**Figure 14:**
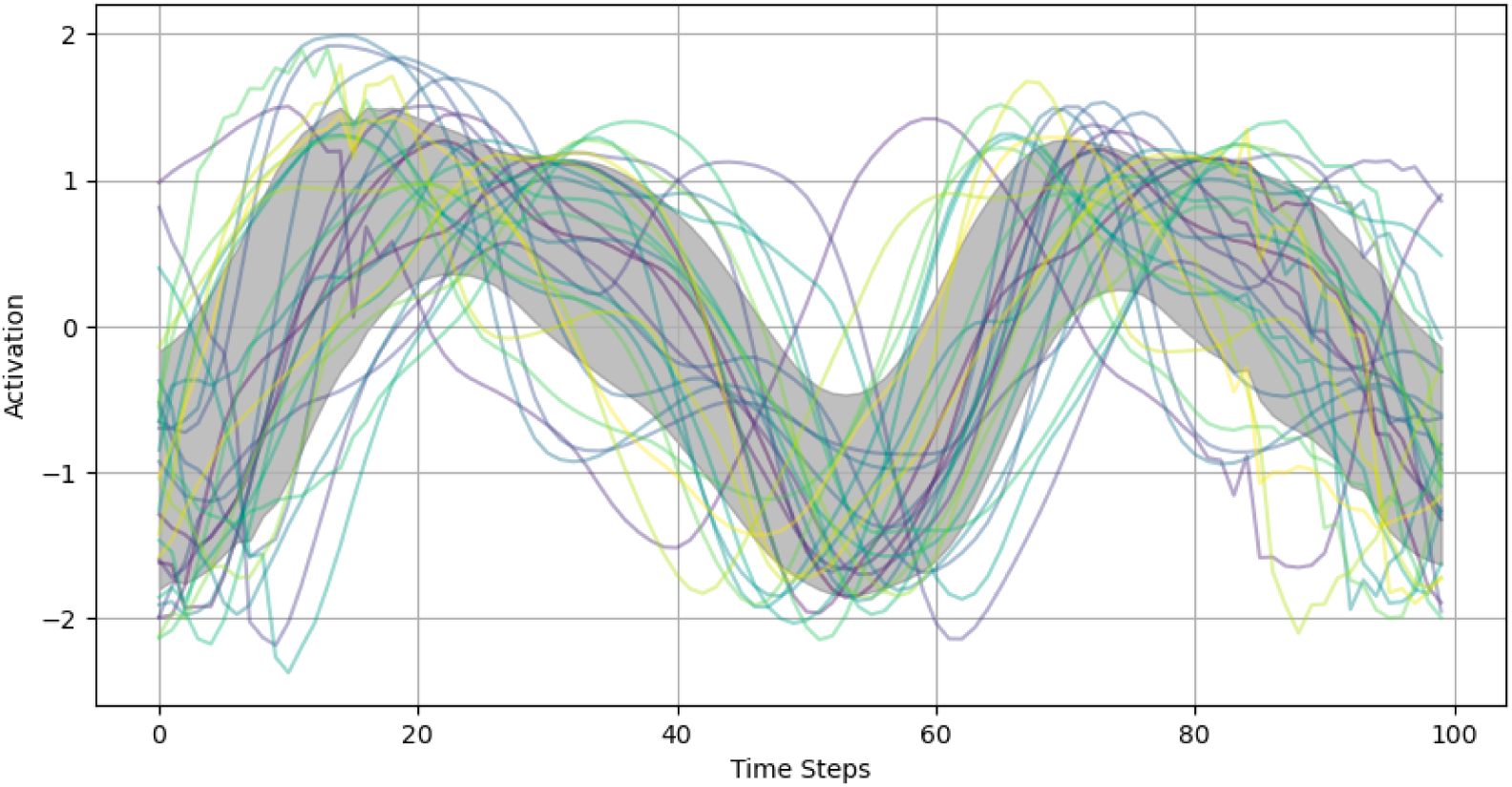
Temporal factors across models.

Overall the analysis employing TCA and hierarchical clustering provided insights into the learning dynamics and neuron behaviour of the simple model trained under identical conditions. Our study identified the variability and consistency in neuron activation across different models, showcasing how the architecture can learn identical tasks using different neuronal assemblies. This research highlights the balance between neuron activity and model accuracy, suggesting that optimal learning may be achieved with a moderate number of highly active neurons, balancing robust training performance with effective generalisation. These findings allow for further explorations into model optimisation, adjustments and further analysis of similar spiking models.

### 7.c. Learning delays

| Section authors | Karim G. Habashy | Balázs Mészáros |
| --- | --- | --- |
| Notebooks | Learning delays, Learning delays (v2), Vanilla sound localization problem with a single delay layer (non-spiking) |  |

This section introduces a simple method to solve the sound localization problem with only learnable delays. Additionally, it discusses a method that learns both weights and delays, introduced in (Hammouamri et al., 2024).

#### 7.c.i. Introduction

Studying the computational properties of axonal transmissions goes as far back as in (Jeffress, 1948). In this study, it was shown that with the right network setup, axonal delays can be utilized to transform a temporal cue to a spatial one for sound localization. This study was a leading study in terms of using delays explicitly to explain a neuronal function. It paved the way for others to follow, like the study by (Kempter et al., 2001), where they investigated the question of how ITD computation maps can arise ontogenetically in the laminar nucleus of the barn owl. They showed that interaural time differences (ITD) computational maps emerge from the combined effect of a Hebbian spike-based learning rule and its transmission along the presynaptic axon. Thus, from this study, another role of axonal delays can be inferred.

They shape network structure when coupled with temporal learning rules. Based on this insight, several studies investigated the combined effect of spike timing-dependent plasticity (STDP), axonal conduction delays and oscillatory activity on recurrent connections in spiking networks. (Kerr et al., 2013) demonstrated the selective potentiation of recurrent connections when the beforementioned computational considerations are taken into account. Also, (Kato & Ikeguchi, 2016) showed that neural selection for memory formation depends on neural competition. In turn, for neural competition to emerge in recurrent networks, spontaneously induced neural oscillation coupled with STDP and axonal delays are a perquisite.

Coupling conduction delays with STDP seems like a reasonable choice. The sign of the STDP rule depends on the order of post- and pre-synpatic spiking, which axonal delays can effectively reverse. For example, if the presynaptic spikes arrive at the synapse before the backpropagated action potential this would lead a synaptic depression. However, reducing the axonal transmission speed would lead to potentiation. In this line of thought, (Asl et al., 2017) studied the combined role of delays and STDP on the emergent synaptic structure in neural networks. It was shown that, qualitatively different connectivity patterns arise due to the interplay between axonal and dendritic delays, as the synapse and cell body can have different temporal spike order.

Aside from their role in modeling cortical functions or shaping a network’s synaptic structure, another line of research emerged from the seminal work by (Izhikevich, 2006). They showed that when including conduction delays and spike-timing dependent plasticity (STDP) into their simulation of realistic neural models, polychronous groups of neurons emerge. These groups show time-locked spiking pattern with millisecond precision. Subsequent studies investigated the properties and functions of such neuronal groups. For example, (Szatmáry & Izhikevich, 2010) demonstrated the natural emergence of large memory content and working memory when the neuronal model exploits temporal codes. Specifically, short term plasticity can briefly strengthen the synapses of specific polychronous neuronal groups (PNG) resulting in an enchantment in their spontaneous reactivation rates. In a qualitatively different study, (Eguchi et al., 2018) showed that networks that exhibit PNG possess potential capabilities that might solve the dynamic binding problem. These networks respond with stable spatio-temporal spike trains when presented with input images in the form of randomized Poisson spike trains. The functionality of these kind of networks emerged due to the interplay of various factors including: i) random distribution of axonal delays ii) STDP ii) lateral, bottom-up and top-down synaptic connections.

Finally, it should be noted that most of the studies that incorporate axonal and/or dendritic delays, include them as a non-learnable parameter. Few studies investigated the possibility of training transmission delays in order to enhance the computational capabilities of spiking neural networks (SNN). (Matsubara, 2017) proposed an algorithm that modifies the axonal delays and synaptic efficacy in both supervised and unsupervised approaches. The learning method used approximates the Expectation-Maximization (EM) algorithm and after training, the network learns to map spatio-temporal input-output spike patterns. Thus, EM is one way to train SNN that are cast as probabilistic models. Another approach that exploits the massive infrastructure that is laid out the deep learning literature is the work by (Hammouamri et al., 2024). In this work, delays are represented as 1D convolutions through time, where the kernels include a single per-synapse non-zero weight. The temporal position of these non-zero weights corresponds to the desired delays. The proposed method co-trains weights and delays and is based on the Dilated Convolution with Learnable Spacings (DCLS) algorithm [(Khalfaoui-Hassani et al., 2023)].

In this work we propose a delay learning algorithm that is simple and efficient. The delay learning is mediated by a differentiable delay layer (DDL). This layer can be inserted between any two layers in an SNN in order to learn the appropriate delay to solve a machine learning task. This DDL is architecture agnostic. Also, the method is designed to learn delays separately from synaptic weights.

#### 7.c.ii. Methods

The DDL is, mainly, based on a 1D version of the spatial transformer (STN) network (Jaderberg et al., 2015). The STN is a differentiable module that can be added into convolutional neural networks (CNNs) architectures to empower them with the ability to spatially transform feature maps in a differentiable way. This addition leads to CNNs models that are invariant to various spatial transformations like translation, scaling and rotation.

Image manipulations are inherently not differentiable, because pixels are a discrete. However, this problem is overcome by the application of an interpolation (for example bi-linear) after the spatial transformation.

The DDL is a 1D version of the spatial transformer where the only transformation done is translation. Translation of a spike along the time dimension can be thought of as a translation of a pixel along the spatial coordinates. The general affine transformation matrix for the 2D case takes the form in the following equation:

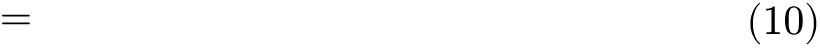

In the above equation, 𝑠*r*_1_, ∼𝑠*r*_2_, ∼𝑠*r*_3_, ∼𝑠*r*_4_ are the elements responsible for the linear transformations of scaling and rotation. *t*_*x*_∼𝑎𝑛𝑑∼*t*_𝑦_ are the translations in the x-axis and y-axis respectively. *x*_*t*_∼𝑎𝑛𝑑∼𝑦_*t*_ are the location of a spike/pixel (in case of spikes y = 0) in the target/output grid, while *x*_𝑠_∼𝑎𝑛𝑑∼𝑦_𝑠_ are the location of the source grid. A grid can be an image or a 2D array of spike trains. For the case of only translation along the x-axis, the affine transformation matrix becomes:

Conventionally, for the spatial transformer, after the projection of the target grid onto the source grid comes the process of interpolation as the transformed pixel might not coincide with a representative one in the source grid/image. Hence, interpolation is performed to estimate the value of the transformed pixel from the surrounding ones. However, applying this process to spike trains can lead to a distortion of the spikes as the allowed values are only 1s and 0s. To avoid this, the translation element *t*_*x*_ should be multiples of the minimum delay. Thus, any transformed spike location from the target grid will find a matching spike location in the source grid. Interpolation (for example bi-linear) will pick the coincident source spike with a weighting of one and provide zero weighting for any other nearby spikes. This process is summarized visually in Figure 15. The input spike trains to the DDL should be padded by zeros so as to not lose information after translation.

**Figure 15:**
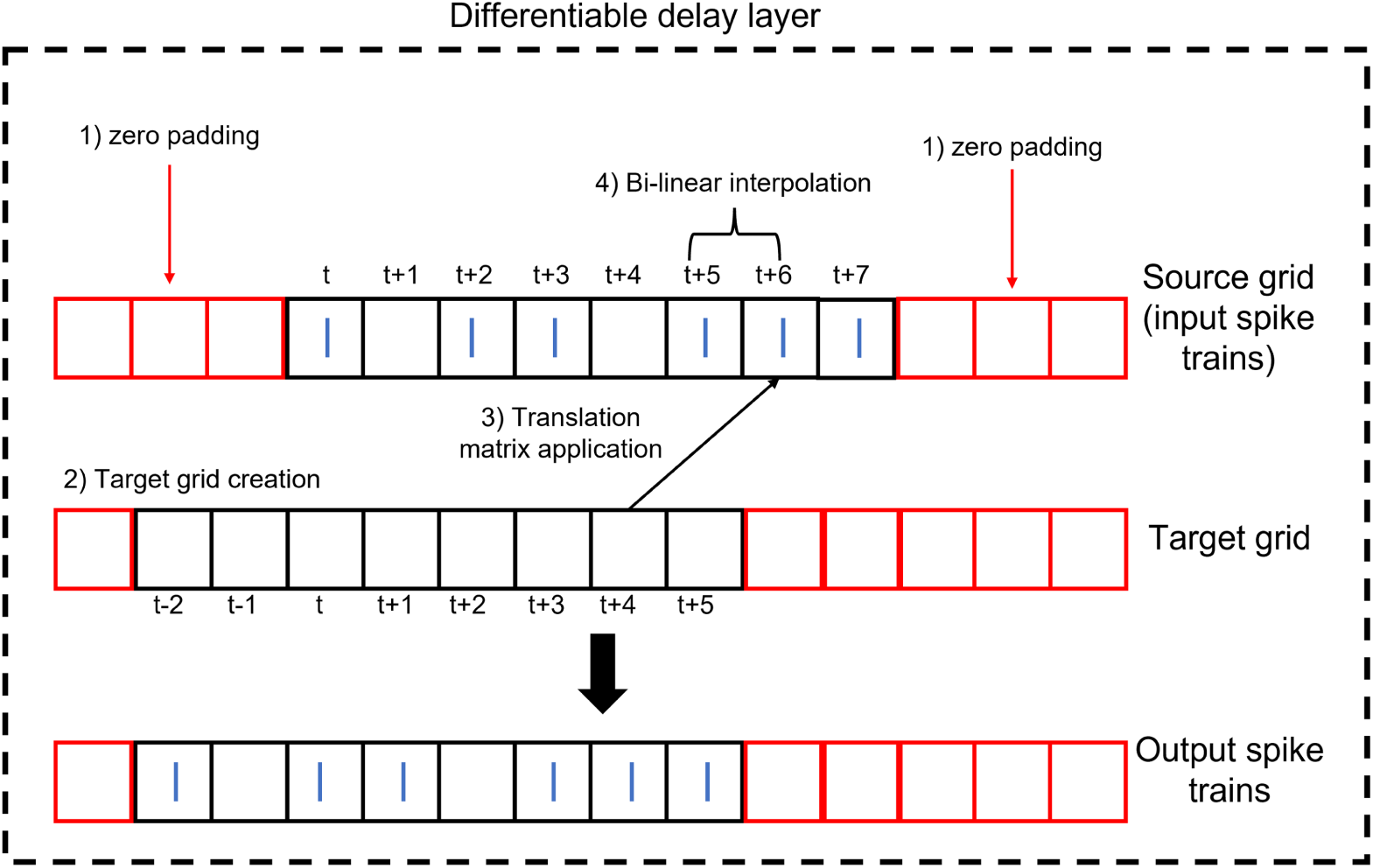
Structure of the DDL. The DDL shifts an input spike train by applying translation then interpolation.

Only the DDL is needed to solve the sound localization problem, where the output classes are the target IPD. As shown in Figure 16, the DDL inserted between the input and output nodes is sufficient to solve the sound localization problem. In this cases, the weights are set to one and the biases to zero.

**Figure 16:**
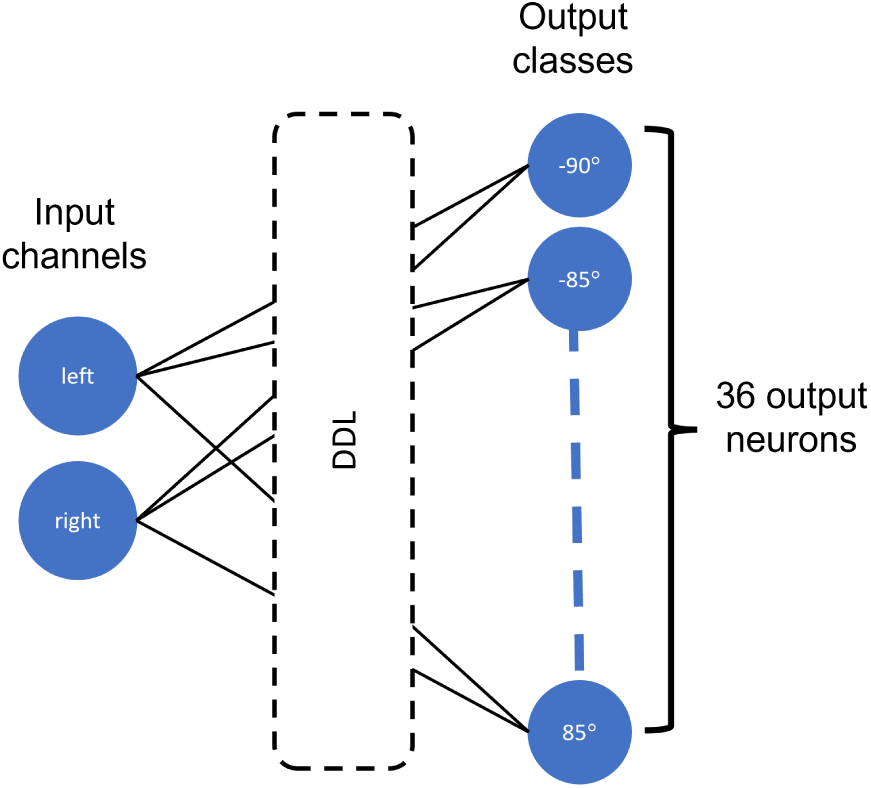
The model architecture. The DLL inserted between the input and out nodes is sufficient to solve the sound localization problem. The output nodes are IPD classes spanning the range [−90^𝑜^, ∼85^𝑜^]∼𝑖𝑛∼5^𝑜^ steps.

Though the DLL shifts the spike trains in a differentiable way, this shift has a meaningful impact through a multiplicative synapse. Also, to facilitate learning, a non-spiking output was utilized. Thus, these two points taken together, the voltage at the output takes the form:

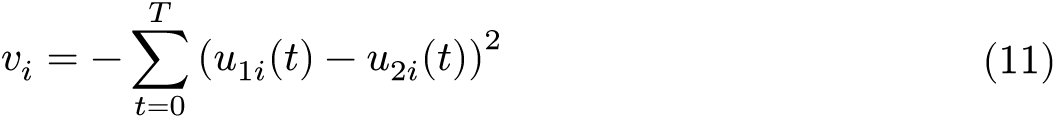

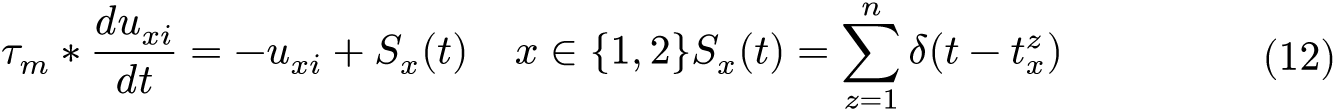

𝑣_𝑖_ is the voltage at output node 𝑖.∼𝑢_*x*_𝑖 is the dendritic potential. 𝜏_*m*_ is the time constant of a dendritic branch. 𝑆_*x*_(*t*) is the input spike train. The form (𝑢_1𝑖_(*t*) − 𝑢_2𝑖_(*t*))^2^ looks like a squared error and might, at first glance, seem not biological, but the expanded term (as seen from the following equation) is a from of multi-synaptic multiplicative-additive interaction.

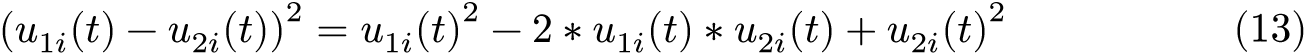

The synaptic delay learning method, employing Dilated Convolutions with Learnable Spacings, operates by delaying spike trains through a 1D convolution featuring a single non-zero element, equivalent to the synaptic weight, positioned at the appropriate delay value.

This method also uses interpolation to identify the optimal delay, facilitating the learning of delays with weights through backpropagation through time in arbitrarily deep SNNs. As we have implemented the method precisely as described in the original paper (with the exception of hyperparameters), we direct the reader to the original paper for a comprehensive understanding(Hammouamri et al., 2024).

#### 7.c.iii. Results and discussion

In this section, the problem complexity is increased up to 36 output units spanning an IPD range of [−90, ∼85] with a step of 5^𝑜^. Employing the DDL to solve such a task leads to the spike raster plots shown in Figure 17.

**Figure 17:**
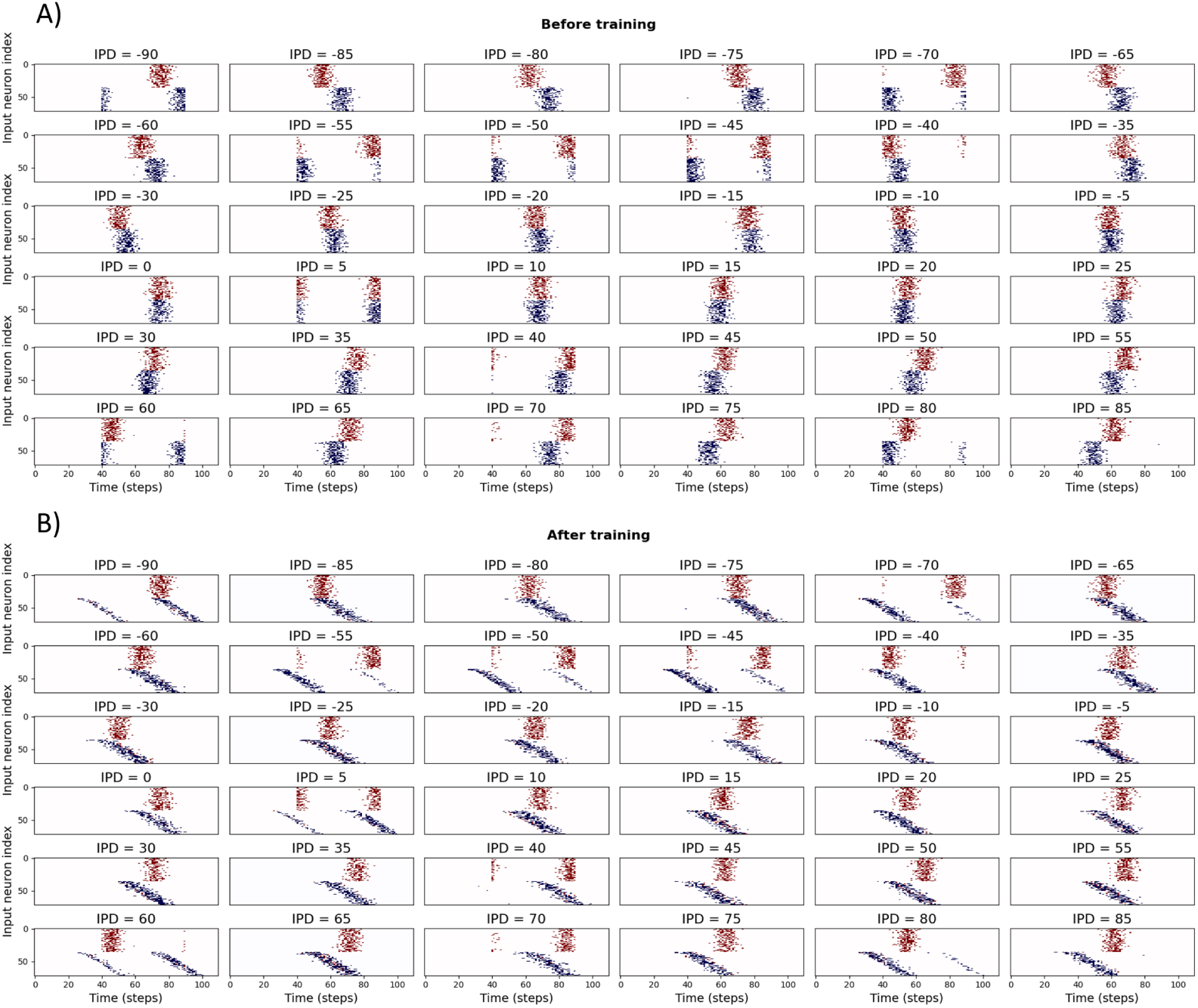
Spike histograms for IPDs before and after training.

To facilitate the search for a solution while using the DDL, we assume that the delay lines coming from one ear is fixed, while the delay lines from the other ear is adaptable starting from an initial default value. Thus, the learned delays can be thought of as the change in delays, which is added to the default delays. This approach leads to the results shown in Figure 17B, where the top of the spike histograms is fixed, while the bottom half has a graded shift in its spike plots. Such results were achieved with the application of time constant decay that leads to a decrease in loss as shown in Figure 18A.

**Figure 18:**
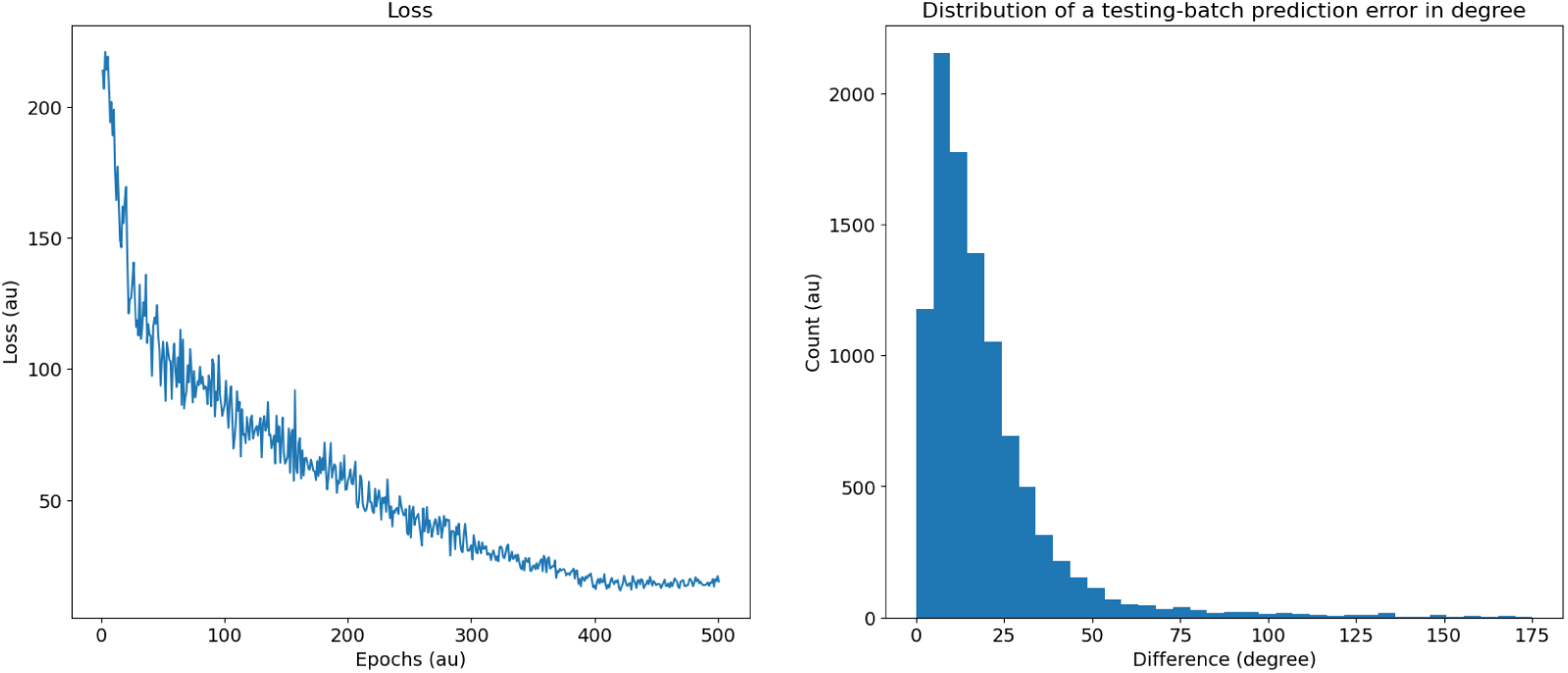
Performance metrics one. A) Loss as a function of the epochs. B) The difference between the True and predicted IPDs in a test batch.

Further analysis of such solutions warrants testing them in different forms. In this regards, we start by displaying the distribution of the errors between the true IPDs and predicted IPDs in a test batch as shown in Figure 18B, which shows an almost log-normal distribution. In addition, we show the confusion matrix for both the training and testing batches on the right images of Figure 19, and the difference between the true and predicted IPDs on the left images of Figure 19.

**Figure 19:**
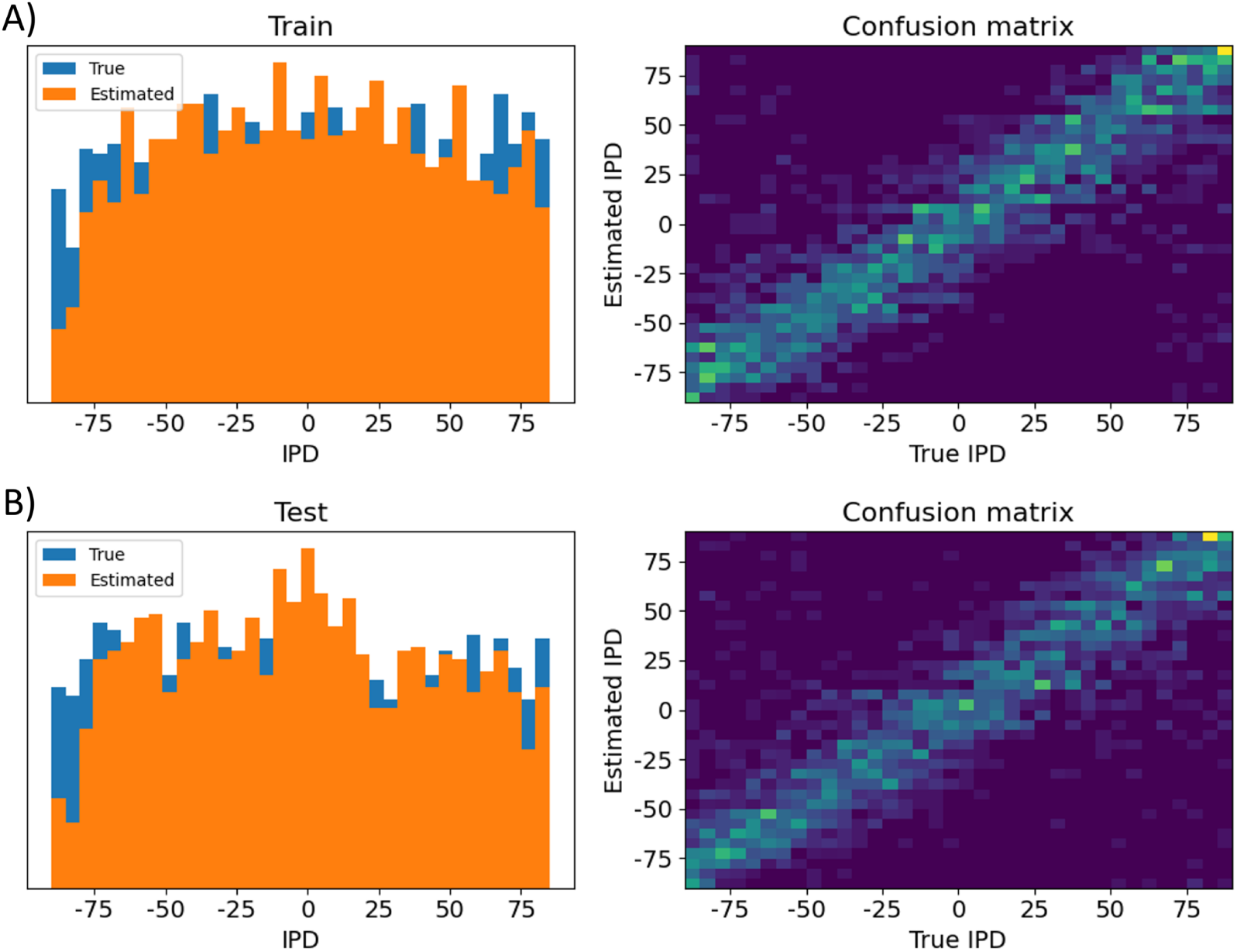
Performance metrics two. Here is shown the distribution and confusion matrix between true IPD values and A) training batch IPD estimates, B) testing batch IPD estimates.

Similar to the other cases, the DCLS architecture is trained for classification across 12 classes, as shown in Figure 20. While delays were manually added to the data in other cases, it is not feasible to ascertain if the method learns identical delay values due to the implementation of varied conduction delays for each synapse. However, in terms of performance, similar accuracy is achieved.

**Figure 20:**
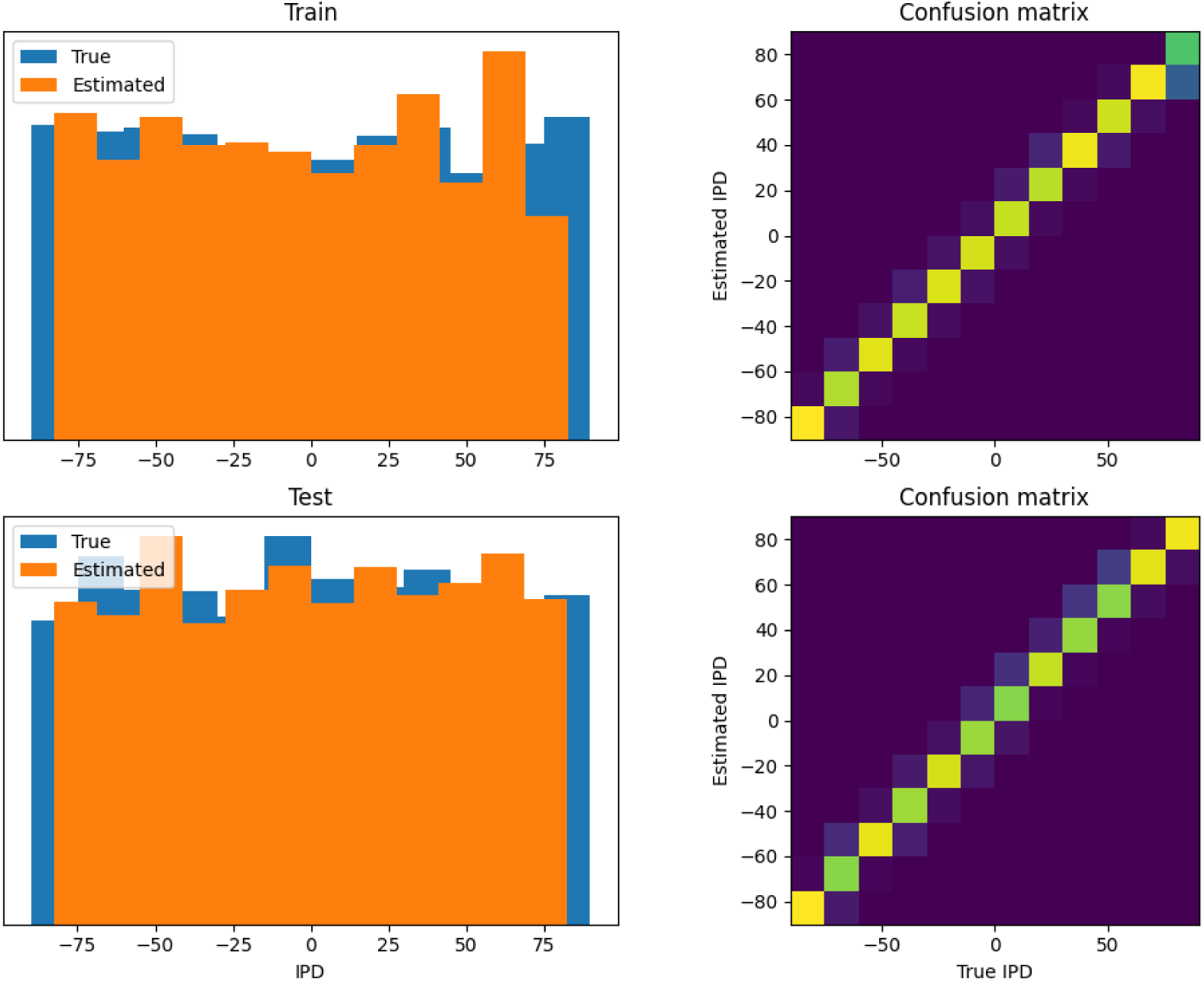
The same performance metrics for DCLS

Learning synaptic delays with weights enables the visualization of the ‘receptive field’ of postsynaptic neurons, as illustrated in Figure 21. Five randomly chosen neurons from the hidden layer are plotted, revealing clear spatiotemporal separation of excitation and inhibition.

**Figure 21:**
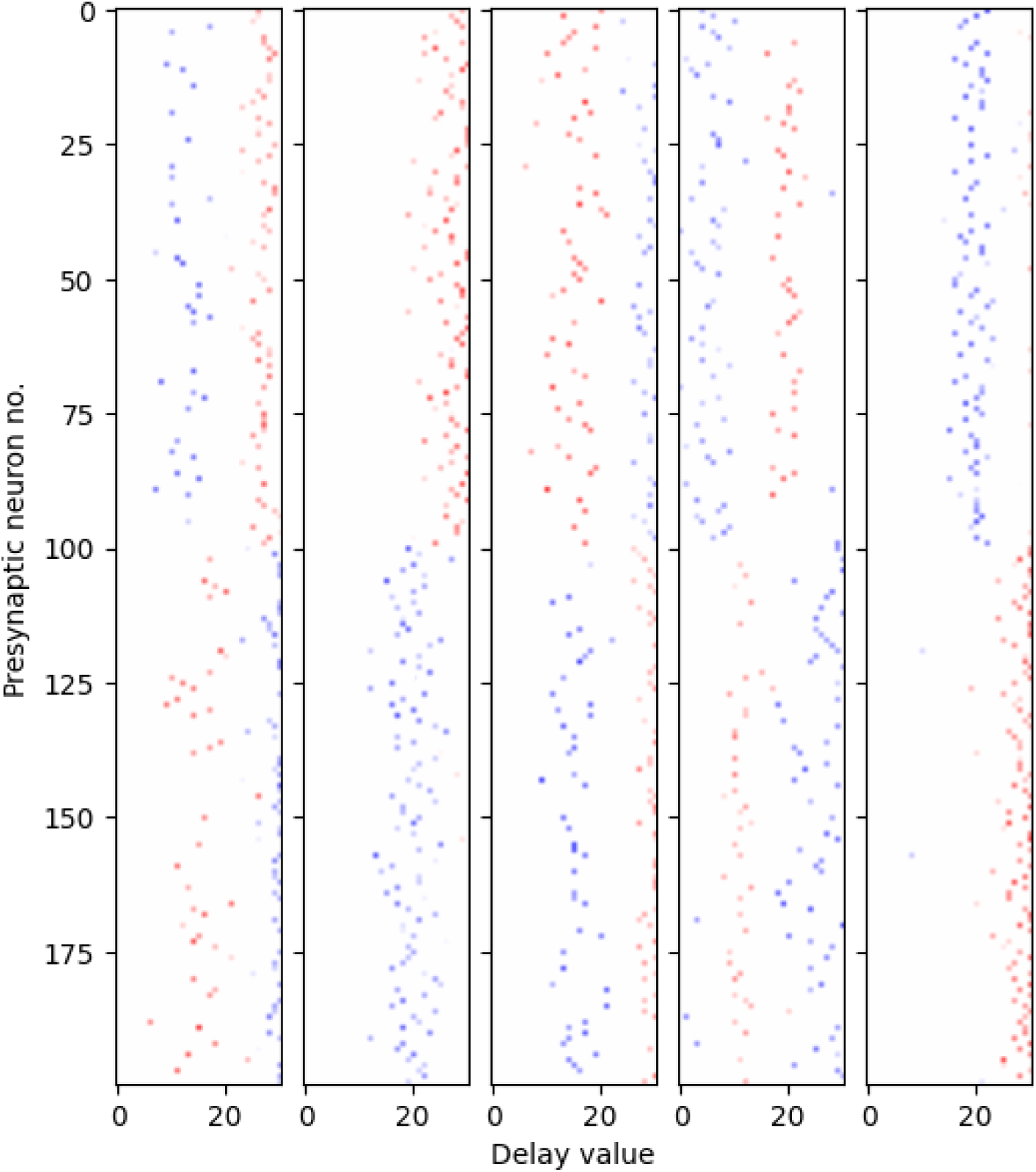
Receptive fields of 5 randomly chosen postsynaptic neurons. The x-axis represents the presynaptic neuron index, while the y-axis displays the learned delay value. Colors indicate the sign of the weight (excitation or inhibition), with transparency denoting magnitude.

### 7.d. Contralateral glycinergic inhibition as a key factor in creating ITD sensitivity

|  |  |
| --- | --- |
| Section authors | Francesco De Santis, Alberto Antonietti |
| Notebooks | Inhibition Model Notebook |

#### 7.d.i. Highlights

Here we use a more biologically inspired model to overcome some limitations that have been highlighted in the classic Jeffress model, whose existence, in mammals, is still debated. In fact, axonal delay lines have not been found in the mammalian MSO. We focused then our attention on the inhibitory inputs to the MSO, which were found both anatomically (there are inhibitory ipsilateral and contralateral pathways) and computationally (in our model) to have a central role for coding ITDs. Experiments with inhibition blocked (by a glycine antagonist: strychnine) shown indeed a loss of ITD-coding in the MSO.

#### 7.d.ii. Introduction

For the vast majority of animals, sound localization is realized through two classes of hidden acoustic cues: binaural and spectral cues. In humans, binaural cues alone are sufficient to discriminate azimuth angles between −90° to +90° degrees, whereas psychoacoustic evidence has shown how spectral cues have a role in the solution of front-back ambiguities and in angle recognition along the vertical plane (i.e. elevation angles) (Gelfand, 2010). Focusing on binaural cues, the two main signals exploited by different animals in creating an acoustic spatiality are interaural time and level differences (ITDs and ILDs). Their role and their dominance over each other depend mainly on animal head size and hearing range in terms of frequencies.

It is clear at first glance how ITDs are longer for species with large head sizes, conversely, these are much shorter for smaller animals, and therefore harder to be neuronally encoded. ILDs on the other hand occur when the wavelength of the sound stimulus is shorter than the size of the animal’s head, which then generates a significant attenuation at the contralateral ear. For this reason, ILDs’ significance with respect to ITDs’ is greater in smaller animals with a middle ear specialised for the transmission of high frequencies (thus having a high-frequency hearing range). This was the case with early mammals, which in fact only developed structures dedicated to processing ILDs.

Even today, many small mammals rely exclusively on ILDs to perform sound localization and the main neural structure involved in the processing of this cue is the Lateral Superior Olive (LSO), a nucleus placed within the superior olivary complex in the mammalian brainstem. Only afterwards the selective pressure led some mammalian species to evolve a second specialized structure, the Medial Superior Olive (MSO), for the processing of ITDs solely. This was probably due to an increase in body size that provided production of low-frequency communication calls and thus the need to implement coding of ITDs as well (Grothe & Pecka, 2014). Humans, whose hearing range is shifted towards lower frequencies with respect to many mammals, possess both a large LSO and a well-developed MSO. As a matter of fact, psychoacoustic evidence has shown how both ITDs and ILDs are used for horizontal sound localization. In particular, the MSO can be considered as a refined processing stage with respect to the LSO for the localization of low-frequency sounds, at which ILD cues are not available (Grothe & Pecka, 2014).

Therefore, it is important to outline the coding strategy for ILDs in the LSO to fully grasp the development of ITD sensitivity in the mammalian MSO. LSO principal cells have a bipolar dendritic tree that receives excitatory input from the ipsilateral ear and inhibitory input from the contralateral one. All mammals appear to apply one common neural strategy for processing ILDs, which consists of subtraction between these two inputs. For instance, neurons in the right LSO will be more excited when sound arrives from the right and more inhibited when sound arrives from the left. The function that describes LSO neurons’ firing rate according to different azimuth angles is usually a sigmoid, presenting the highest values for sound coming from the hemispace ipsilateral to the nucleus, with a steep slope centred on frontal angles (i.e., close to 0°), suggesting a rate-coding strategy for identifying different ILD values. Although the on-off nature of this subtraction strategy, for which exquisite timing of inhibitory influences appears not to be a key prerequisite, the two major inputs to the LSO are specialized for high-fidelity temporal transmission. An explanation is found in the fact that the subtraction process happening in LSO principal cells is realized in a phase-locked way with respect to the stimulus. This implies a purely suppressive coincidence mechanism happening at each period of the phase-locked inputs to the LSO (spiking occurs unless binaural coincidence is happening).

The MSO, on the other hand, receives two additional inputs compared to the LSO: a contralateral excitation and an ipsilateral inhibition. As a result of experimental observations, the main hypothesis is that the combination of these four inputs has converted the suppressive coincidence mechanism present in the LSO into an excitatory coincidence one for the detection of ITDs in the MSO (spiking occurs only if binaural coincidence is happening) (Grothe & Pecka, 2014).

Nevertheless, the most famous model that attempts to explain the functioning of the MSO, namely the Jeffress model, considers only the two excitatory inputs arriving at the MSO, assuming an array of labelled neurons providing binaural coincidence detection for specific values of ITDs, due to the presence of axonal delay lines that can create this type of sensitivity (Jeffress, 1948). For many years, Jeffress’s remained the most accepted proposal, given the discovery of axonal lines potentially similar to those theorised by Jeffress in the Nucleus Laminaris (NL) of different bird species. However, the presence of similar delay lines in the mammalian MSO, analogous to the avian NL, has never been demonstrated.

The first steps toward a better understanding of the MSO functioning were made through the work of Brand et al. (2002) in which the analysis of *in vivo* recordings from the MSO of the *Mongolian gerbil* showed how all the 20 neurons tested responded maximally to sounds leading in time at the contralateral ear. Peaks in the firing rate of these neurons were found also for (artificial) ITD values higher than the highest possible ones generated by the gerbil head, which correspond to almost 120 *µ*𝑠 for a sound coming at 90° from the contralateral hemispace. This observation led to the hypothesis that peaks in these neurons’ firing rate were not coding for a specific ITD value in gerbil-relevant physiological space, suggesting instead a mapping made by the slopes in the activity curves (McAlpine & Grothe, 2003), in a way similar to the rate-coding strategy adopted for ILDs in the LSO (even if in LSO cells the slope is positive passing from contralateral to ipsilateral sound sources, oppositely from what happens in MSO cells response). The idea of a topographic map at the level of the MSO portrayed by different peaks in cell activity (i.e., peak-coding) risks then to be discarded in favour of a less refined rate-coding strategy, in which recognition of the ITD value generated by an input sound should occur at a higher level (e.g., at the Inferior Colliculus).

Brand et al. (2002) and Pecka et al. (2008) explored also the physical mechanisms underlying the shift in peaks’ activity towards contralateral ITD values. Since the presence of axonal delay lines had been discarded, there had to be another neural mechanism capable of compensating for the external ITD value, causing a growth in the activity of MSO neurons for gradually more and more contralateral sounds. The results show how the two inhibitory inputs to the MSO, which were not considered in the Jeffres model, had instead a central role in this process. By blocking the glycinergic inhibition to the MSO neurons by means of the application of its antagonist, strychnine, it was observed the loss of the peak shift in all their response activity. Having now a symmetry in all the neuron responses, with a peak in the activity corresponding to null ITD values (i.e., 0° azimuth angle), all the information present in the MSO responses had been lost. Inhibition has thus a central role in identifying the ITD values in the MSO, whether the coding strategy adopted consists of peak-coding or rate-coding. However, even in this case, although numerous proposals have been made, it is still unclear how the two inhibitory inputs manage to cause this higher excitability of MSO neurons for contralateral sounds and thus the peak shift observed experimentally. Some experimental studies have reported how inhibitory inputs, especially the contralateral one, can arrive slightly in advance of the excitatory inputs from the corresponding sides, thus influencing the way ipsilateral and contralateral inputs add up in the MSO neurons and thereby generating a maximum response for contralateral ITD values (Myoga et al., 2014; Roberts et al., 2013). Nevertheless, other work has characterized the hypothesis of anticipation of inhibitory inputs as implausible (vanderHeijden et al., 2013). For this reason, in our work, we explored the validity of another hypothesis, proposed by Myoga et al. (2014), for which the neural mechanism employed by the MSO relies on the shape of post-synaptic potentials (PSPs), both excitatory (EPSPs) and inhibitory (IPSPs). Therefore, we developed a realistic *in silico* model of the mammalian brainstem and tested different sets of time constants that then govern the shape of these PSPs.

#### 7.d.iii. Methods

Inspired by the neurophysiological data, we implemented a complex spiking neural network in Python using the NEST Simulator framework (Spreizer et al., 2022). The different neuronal populations composing the brainstem circuit and their interconnections are depicted in Figure 22.

**Figure 22:**
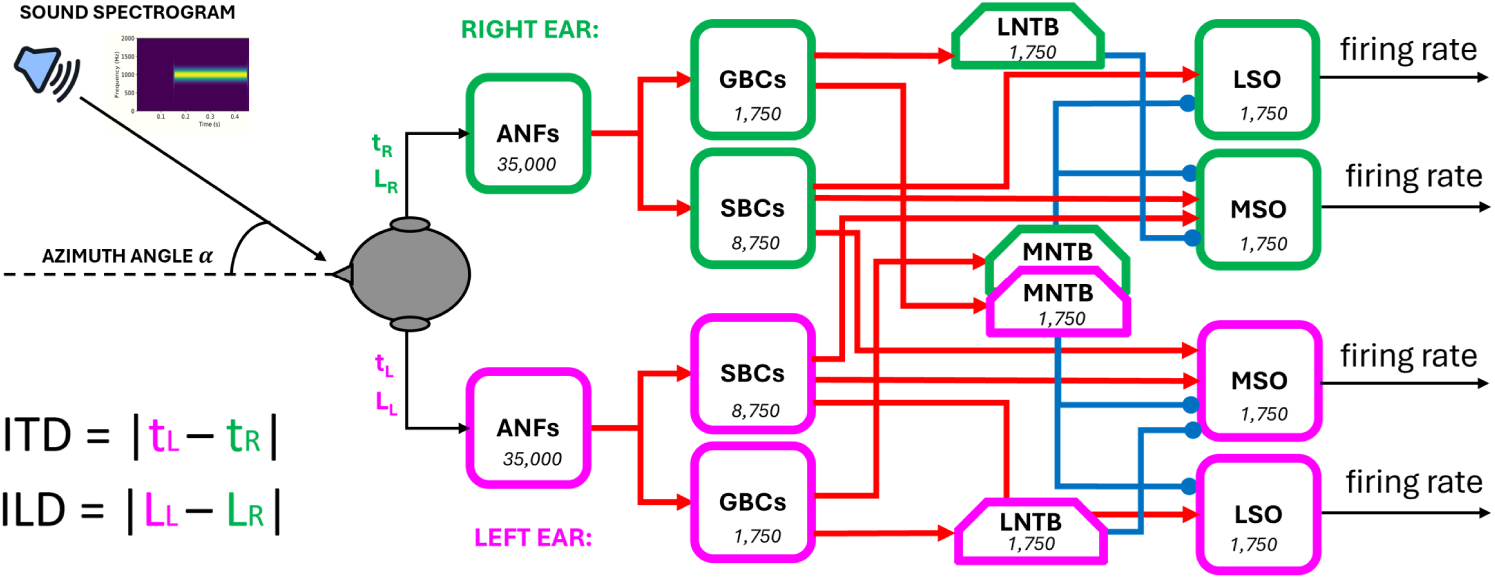
The end-to-end brainstem model with the network diagram. The number of neurons for each population is written in the respective block

The principal inputs to the network are the spectrogram of a sound stimulus arriving at both ears and the azimuth angle from which univocal values of ITD and ILD are computed. With regards to the spectrogram, we covered the whole human audible range of sound between a minimum frequency of 20 Hz and a maximum of 20 kHz. We subdivided it into 3500 channels since this value is the most likely estimate of the number of inner hair cells (IHCs) present in the human cochlea. With the intent to mimic the physiological distribution of IHCs along the basilar membrane, the width of each frequency channel was not constant throughout the range but grew exponentially (Fettiplace, 2023).

For modelling of the auditory nerve fibres (ANFs), we used the `pulse_packet_generator`, a built-in NEST device that produces a spike train containing Gaussian spike clusters centred around given times. These devices mimic the actual behaviour of ANFs subjected to a pure tonal stimulus, which, like all other neuronal populations involved in sound processing, manifest a phase-locked response. The firing of these neurons tends to occur only at a certain restricted phase of the incoming sine wave of sound, due to the working principle of the inner hair cells placed in the cochlea (Yin et al., 2019). `pulse_packet_generator` allowed us to define the number of spikes in each packet, with spike times that are normally distributed with respect to the central time of the pulse. This allowed us to implement ILDs by varying the `spikes_number` parameter according to the input azimuth angle. A source of noise was introduced by setting the standard deviation of the random displacement from the centre of the pulse equal to 0.1 ms. The value of ITD was instead computed considering the interaural path difference, which is the difference between the paths travelled by the sound starting from the source and arriving at the two ears and dividing it by the speed of sound in air (i.e., 330 m/ s). All other cell populations were implemented in a manner faithful to their biological counterparts, starting with the bushy cells (spherical and globular, SBCs and GBCs) located in the anteroventral part of the cochlear nuclei, proceeding to the glycinergic neurons located in the medial and lateral trapezoidal body (MNTB and LNTB), and finally to the main cells of the medial and the lateral superior olives (MSO and LSO). The MSO principal cells were modelled through the `iaf_cond_beta model`, a simple implementation of a spiking neuron in NEST using integrate-and-fire dynamics with conductance-based synapses. In this model, incoming spike events induce a postsynaptic change of conductance modelled by a beta function. The use of a beta function allowed us to change independently the time constants of rise and fall of the conductance change and thus modify both the excitatory and inhibitory post-synaptic potential shapes in the MSO. In this way, we could explore different sets of values and attempt to validate our hypothesis about how inhibitory inputs can code for different ITD values in MSO cells, see Section 7.d.ii. All the other neuronal populations, except for ANFs and MSO cells, were modelled with the `iaf_cond_alpha` since there was no need for modifying the PSP shape of their inputs.

For the validation of the complete brainstem network, including both LSO and MSO of both sides, sound stimuli with frequencies of 100 Hz and 1 s duration from different spatial positions were tested. Specifically, we gave the model azimuth angles ranging from −90° to +90° with an interval of 15°. The MSO response was tested both in physiological conditions and with blocked inhibitory inputs, as in the experiments of Brand et al. and Pecka et al.

#### 7.d.iv. Results

As for the LSO response in Figure 23, in our results we obtained the desired subtraction process described in Section 7.d.ii: considering for the sake of simplicity the left LSO, when the sound arrives from a source placed at 90° (i.e., right), the right ear receives sounds earlier and more intensely than the left. As the azimuth angle proceeds toward 0° (frontal position), the firing rate of the left LSO increases while maintaining a constant slope. This steep segment is the most informative part of the left LSO response curve, as a high sensitivity to changing input azimuth angles is guaranteed. Once past 0°, the firing rate ceases to increase steadily, and the response flattens out to high-rate values. Here, the ipsilateral (left) excitation dominates due to louder sounds.

**Figure 23:**
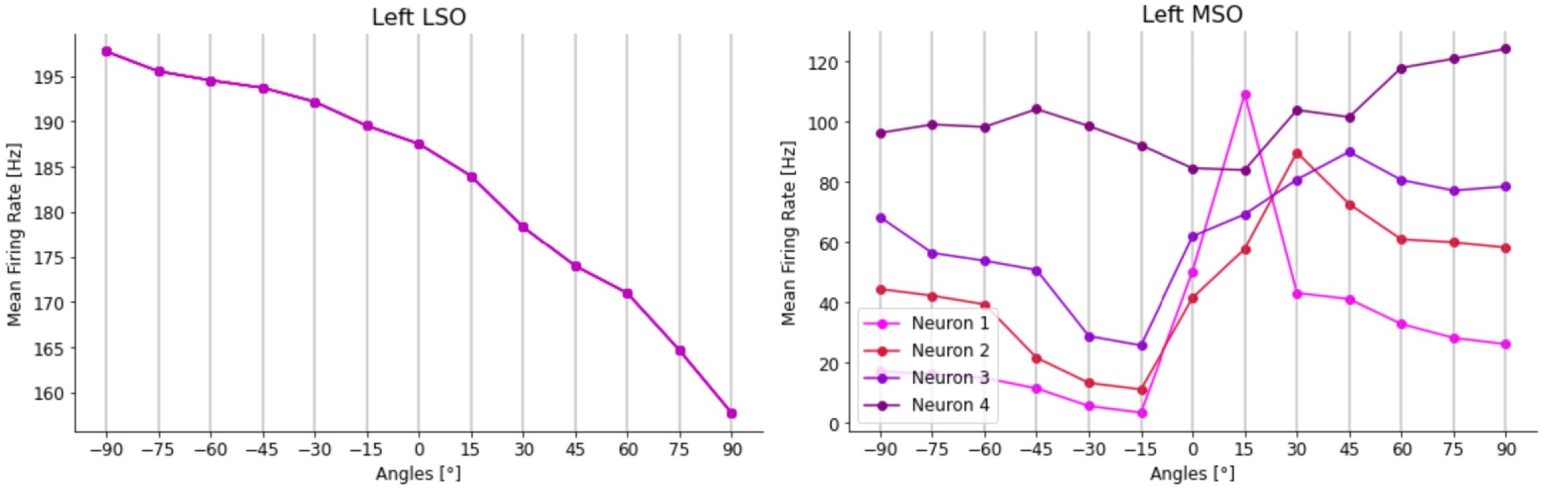
Mean firing-rate responses of the left LSO (on the left) and the left MSO (on the right) after stimulation with a 100 Hz pure tone sound for 1 second at different azimuth angles. For the MSO, four neurons with different time constants are shown. Neuron 1 responds maximally for input at +15°, Neuron 2 at +30°, Neuron 3 at +45°, and Neuron 4 at +90°.

Regarding the MSO instead (Figure 23), the different curves represent the activity of four neurons in the left MSO for which we varied only the value of the time constants of the decay in conductance generated by input spikes for both excitatory (`tau_decay_exc`) and inhibitory (`tau_decay_inh`) inputs. We observed that different combinations of these two values provided coding for a specific angle achieved by a peak at different angles (and thus different ITD values) in that cell’s activity. As observed experimentally, all peaks were present in the contralateral sound space, thus confirming the hypothesis that, in contrast to the coding strategy applied in the LSO, higher activity is present in the MSO for sounds from the contralateral space.

Regarding simulations in which the MSO receives only excitatory inputs (Figure 24), a loss in the coding of different contralateral angles is evidenced by a symmetric firing rate curve, with all the peak values being higher and shifted towards 0° angles with respect to the physiological activity, as measured experimentally in Brand et al. (2002) and Pecka et al. (2008).

**Figure 24:**
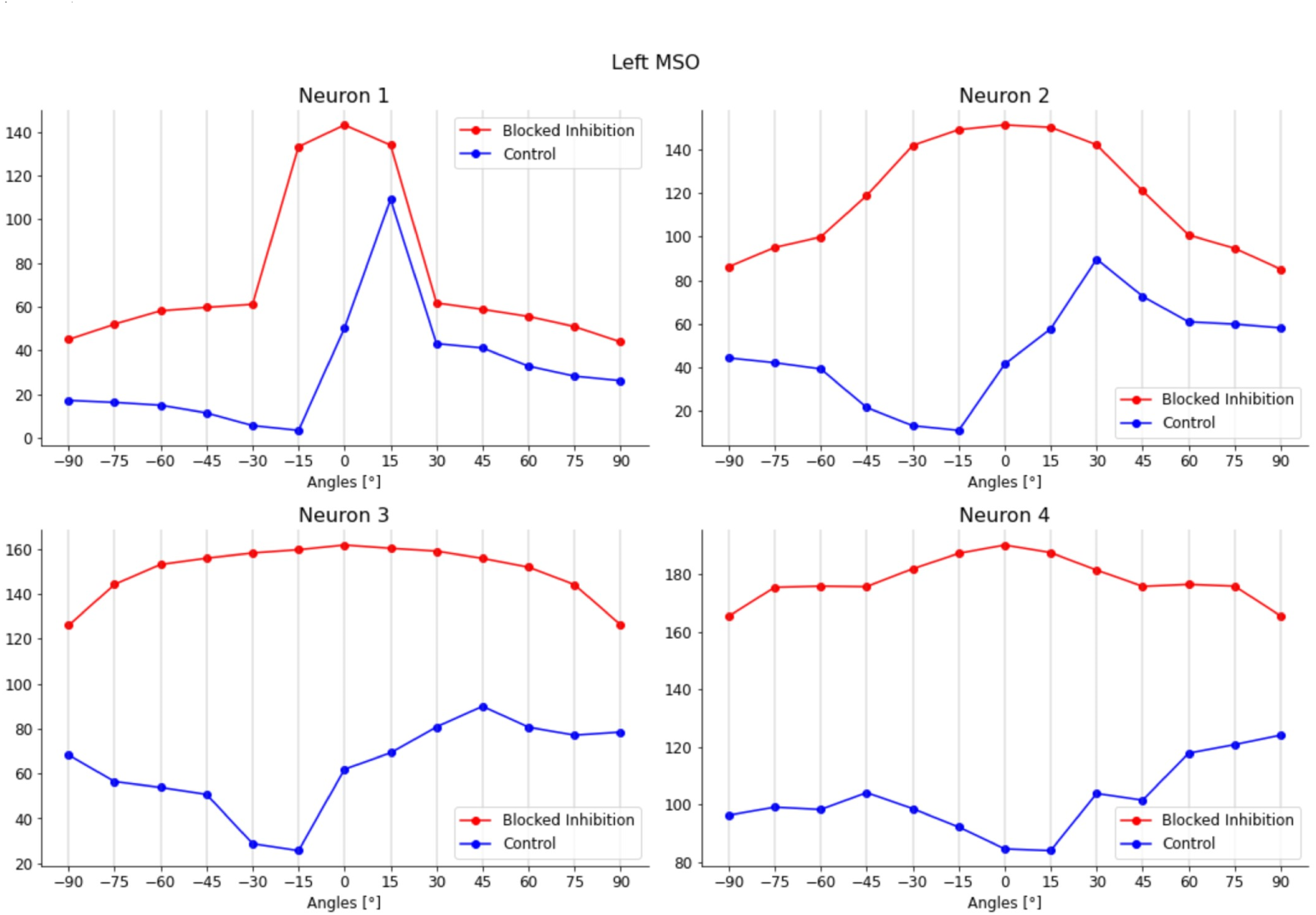
Figure 3: Loss of contralateral ITD peak-coding for the four different neurons in the left MSO: the control condition (with physiological inhibitory inputs) is shown in blue, whereas the curves in red depict the condition with blocked inhibition to the MSO; in the latter scenario, firing-rate values are higher with respect to the former and peaks are shifted to null ITD values, so that the coding of each neuron for a specific azimuth angle is now lost.\

The achievements of this work are to be considered significant in the investigation of the mechanisms used by the mammalian brainstem to perform sound localization. The computational model developed is a good validation platform for the most recent theories concerning the processing of the two binaural cues, ITDs, and ILDs. This model has shown, for pure tones of 100 Hz frequency, the physiological functioning of LSO and MSO. The peak-coding strategy applied for the identification of contralateral angles in each MSO can be considered a refinement of the rate-based localization of sounds happening in the LSO. As described in the Section 7.d.ii, this type of redundancy is also justified by the evolutionary history of spatial hearing mechanisms in mammals. In contrast to birds, where ITDs have always been used as the main binaural cue for sound localization, early mammals first made exclusive use of ILDs as acoustic cues and only later developed a sensitivity to different ITDs, created in tandem with the development of more dedicated low-frequency hearing. This leads to the possible conclusion that strategies similar to those used for the processing of ILDs have also been readapted for the processing of ITDs and that in the brainstem of modern mammals, these two processes occur in parallel, merging at a higher level, and thus providing a more refined and complete spatial map.

## 8. Funding and Acknowledgements

- MG is supported by Schmidt Sciences, LLC. Previously, MG was a Fellow of Paris Region Fellowship Program - supported by the Paris Region, and funding from the European Union’s Horizon 2020 research and innovation program under the Marie Skłodowska-Curie grant agreement No 945298-ParisRegionFP.
- FS and AA work is fully funded by the project “EBRAINS-Italy (European Brain ReseArch INfrastructureS-Italy),” granted by the Italian National Recovery and Resilience Plan (NRRP), M4C2, funded by the European Union –NextGenerationEU (Project IR0000011, CUP B51E22000150006, “EBRAINS-Italy”).
- DFE is funded by the SURF programme (undergraduate research fellowship) from the Simons Foundation (SCGB).
- DFE and BAB acknowledge the use of the UCL Myriad High Performance Computing Facility (Myriad@UCL), and associated support services, in the completion of this work.
- ZF is funded by a NSERC PGS-D Scholarship.
- BM is funded by the be.AI Leverhulme Doctoral Scholarships (Leverhulme Trust).
- UA is supported by the UK Engineering and Physical Sciences Research Council (EPSRC) DTP Studentship 2753922 for the University of Surrey.
- JLR was funded by the Max Plank Society.
- YHL is supported by NSERC PGS-D, FRQNT B2X, and Pearson Fellowship.
- VB is supported by the European Research Council (ERC) under the European Union’s Horizon 2020 research innovation program, grant agreement number 715980.

